# Holistic inference explains human perception of stimulus orientation

**DOI:** 10.1101/2022.06.24.497534

**Authors:** Jiang Mao, Alan A. Stocker

## Abstract

Perception of stimulus features such as orientation is widely considered a Bayesian inference process. In contrast to previous Bayesian observer models, we propose that perception is a *holistic* inference process that operates across all levels of the representational hierarchy. We test this hypothesis in the context of a typical psychophysical matching task in which subjects are asked to estimate the perceived orientation of a test stimulus by adjusting a probe stimulus (method-of-adjustment). We present a holistic matching model that assumes that subjects’ responses reflect an optimal match between the test and the probe stimulus, both in terms of their inferred feature (orientation) but also their higher-level (category) representation. Validation against multiple existing psychophysical datasets demonstrates that the model provides a quantitatively detailed and accurate description of subjects’ response behavior including data that previous models fail to even qualitatively account for.

## Introduction

Perception is considered an inference process that optimally combines noisy sensory signals with prior knowledge about the statistical regularities of the world. Countless studies have argued that models of perceptual inference can be parsimoniously expressed within the probabilistic framework of Bayesian estimation (Knill and Richards, 1996). Perception in this framework equates to finding an optimal estimate of a stimulus feature. A characteristic prediction of Bayesian estimation is that perception is biased towards the peak of the prior density distribution, which has been validated by the results of many perceptual and sensorimotor studies (e.g., Körding and Wolpert, 2004; Stocker and Simoncelli, 2006; Jazayeri and Shadlen, 2010; Kim and Burge, 2018).

A quantitative validation of the Bayesian estimation framework crucially depends on an accurate specification of the prior distribution. Visual orientation is one of the few stimulus features for which the prior distribution is well specified in form of the local orientation statistics in natural visual scenes. These statistics have been repeatedly measured and show robust peaks at cardinal orientations (Coppola et al., 1998; Girshick et al., 2011; Wang et al., 2016). Perceived stimulus orientation is typically biased away from cardinal orientations (De Gardelle et al., 2010; Noel et al., 2021), i.e., it is seemingly “anti-Bayesian” for this natural prior distribution. Recent work has demonstrated, however, that the efficient coding hypothesis (Attneave, 1954; Barlow et al., 1961) provides a powerful constraint on sensory uncertainty to resolve this apparent paradox, leading to a consistent Bayesian interpretation of visual orientation perception (Wei and Stocker, 2012, 2015, 2017). Since then, the Bayesian estimation model constrained by efficient coding (in the following simply referred to as the “Efficient Bayesian estimator”) has demonstrated to offer a unifying account for human behavior in a wide variety of perceptual and working memory tasks (e.g., Taylor and Bays, 2018; Polania et al., 2019; Fritsche et al., 2020; Langlois et al., 2021; Prat-Carrabin and Woodford, 2021).

Despite its promise, however, there are several reasons to question this model’s ability to provide a unifying account of orientation perception. First, a full quantitative validation of the model by comparing the entire predicted response distributions against data is still outstanding; previous studies mainly focused on summary statistics such as estimation bias (but see Taylor and Bays (2018)). Furthermore, there are psychophysical data that are difficult to reconcile with the model. Specifically, Tomassini et al. (2010) reported results of a typical orientation matching experiment where subjects were asked to estimate the orientation of a test stimulus by adjusting a probe stimulus. In half of the trials the stimuli used as test and probe were interchanged. If subjects estimated the orientations of test and probe stimulus independently, one would expect the sign of their estimation biases to flip in those trials; a pair of matched stimuli would yield opposite estimation errors when the assignment of test and probe is interchanged. Tomassini et al. (2010) found, however, that the sign of the bias pattern did not flip in those trials – there was a repulsive bias away from cardinal orientations under both conditions. This result cannot be explained with any model that independently estimates the orientations of the test and the probe stimulus. Last but not least, there is the long-standing notion that higher-level, categorical representations influence perception at the feature level. Several studies have suggested that the perception of visual orientation is affected by a cardinal/oblique category distinction (Rosielle and Cooper, 2001; Wakita,2004). The efficient Bayesian estimator does not provide the possibility to incorporate potential categorical effects unless the orientation prior implicitly reflects the categories, i.e., has peaks at orientations that correspond to the category centers (e.g., Bae et al., 2015). Allowing such an orientation prior, however, generally violates the fundamental Bayesian assumption that the prior distribution reflects the statistical distribution of visual orientations (see above).

Here, we introduce a hierarchical inference model of perception that resolves these issues. What fundamentally separates our proposal from previous models is that we describe perception as a *holistic inference* process, where the percept of a stimulus is jointly represented by the inference outcomes (i.e., the posteriors) at every level of a representational hierarchy. Specifically with regard to orientation perception, we assume that perceived orientation of a stimulus is characterized by inference at both the feature (orientation) as well as potential higher level representations (orientation categories). Our hypothesis reflects the holistic experience we typically associate with perception. Furthermore, the model assumes that cognitive processes downstream of perception (e.g. a decision stage) operate on these holistic perceptual representations of sensory information. We tested our model in the context of a typical psychophysical matching task in which subjects are asked to estimate the perceived orientation of a test stimulus by adjusting a probe stimulus. We show that our model provides a highly accurate account of several existing datasets: the new model not only correctly predicts the non-inverted biases when test and probe stimuli are switched (see above; Tomassini et al., 2010) – something that previous models can not – but also provides a far superior quantitative account for the full error distributions of subjects’ perceptual estimates reported in other experimental studies (De Gardelle et al., 2010; Noel et al., 2021).

## Results

### Holistic perceptual matching

Perception of visual orientation is commonly assessed with a psychophysical matching task (method of adjustment). In this task, a subject is asked to adjust the orientation of a probe stimulus in order to match the perceived orientation of a previously or simultaneously presented test stimulus (Fig. 1a). Typically, the probe stimulus is unambiguous and noise-free leading to the general assumption that the probe orientation is a direct reflection of the subject’s perceptual estimate of the test stimulus orientation, aside from some potential motor noise. Under the noise-free probe condition, the efficient Bayesian estimator has been shown to provide a qualitative accurate account of the repulsive perceptual bias and variance patterns and their dependence on stimulus uncertainty observed in these matching experiments (Wei and Stocker, 2015, 2017; Taylor and Bays, 2018). The model assumes perceived test orientation to depend on a noisy orientation measurement *m* and a prior distribution on orientation *p*(*θ*). Bayesian inference results in a posterior distribution *p*(*θ*|*m*) based on which the perceived orientation (i.e., the optimal estimate) is determined according to a loss function *L_θ_*.

**Figure 1:**
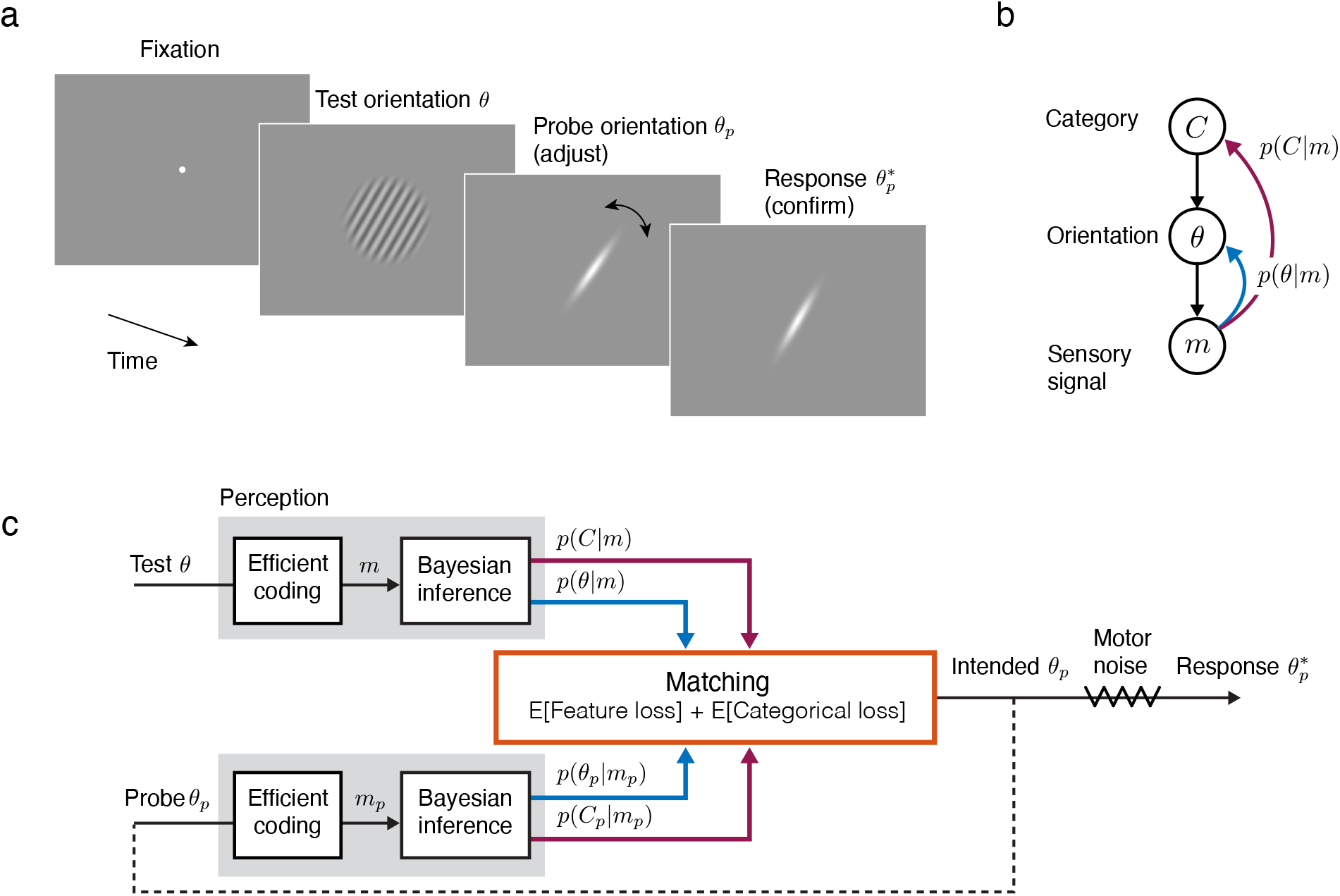
Holistic orientation matching. (a) Typical psychophysical matching task to characterize visual orientation perception. Subjects are presented with a test stimulus with orientation *θ*. Then they are asked to adjust the orientation *θ_p_* of a probe stimulus such that it best matches the perceived test orientation. Subjects typically press a button to confirm their choices, at which time their response 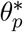 is recorded. (b) Graphical model representing the hierarchical generative process by which a stimulus with orientation *θ* and a higher-level, categorical identity *C* (cardinal/oblique) generates a noisy sensory signal *m*. We assume that perceptual inference is holistic and consists of computing both the posteriors over orientation *p*(*θ*|*m*) (blue arrow) and category identity *p*(*C*|*m*) (purple arrow). (c) Holistic matching model. The model assumes that both the test *θ* and the probe *θ_p_* orientation are efficiently encoded according to the orientation prior *p*(*θ*) (Wei and Stocker, 2015), resulting in sensory measurements *m* and *m_p_*, respectively. As illustrated in (b), perceptual (Bayesian) inference results in posteriors *p*(*θ*|*m*), *p*(*C*|*m*) and *p*(*θ_p_*|*m_p_*), *p*(*C_p_*|*m_p_*), respectively. By minimizing a combined objective that quantifies mismatch at both the feature and category level (Eq. (1)), the model computes the probe orientation that optimally matches the test orientation. Note, that a non-holistic version of the proposed model (i.e., removing the categorical inference pathway - purple arrows) is equivalent to the efficient Bayesian estimator when the probe stimulus is noise-free.

In contrast, the proposed holistic matching model assumes a hierarchical generative process where each stimulus orientation *θ* is associated with a category *C* distinguishing cardinal and oblique orientations (Fig. 1b). Furthermore, we assume that the perceptual inference process consists of inferences at both levels of the hierarchy. The outcome is thus a holistic representation of the perceived orientation stimulus, jointly represented by the posteriors at both the orientation and the category level, *p*(*θ*|*m*) and *p*(*C*|*m*), respectively. The key innovation is that the matching stage operates on these holistic perceptual representations. That is, the model assumes that the observer aims to adjust the probe orientation *θ_p_* until the percepts of the probe and the test stimulus optimally match at both representational levels (Fig. 1c). We express this as finding the probe orientation that minimizes a weighted average of the expected mismatch at the orientation (*L_θ_*) and the category level (*L_c_*), thus

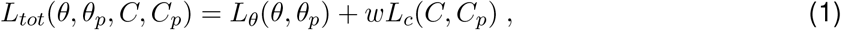

where *w* > 0 is the relative contribution of the categorical mismatch. We define the mismatch at the orientation level *L_θ_* as the cosine difference between the test and the probe orientation, while the mismatch at the categorical level *L_c_* is assumed to be a constant penalty if test and probe belong to different orientation categories and zero otherwise. Finally, we assume that subjects’ responses 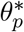 represent noisy samples of the optimally matching probe orientation *θ_p_* due to additive, constant motor noise (*Methods*).

Simulations shown in Fig. 2 illustrate how and why the predictions of the holistic matching model differ from those of the efficient Bayesian estimator. We assume the prior distribution of visual orientation *p*(*θ*) to reflect the orientation statistics in natural scenes. Previous studies have shown that these statistics are relatively robust with regard to the specific methods they were measured with and the image content of the natural scenes they were computed for, showing characteristic peaks at both cardinal orientations (Coppola et al., 1998; Girshick et al., 2011; Wang et al., 2016). However, outdoor scenes containing fewer man-made objects typically show less pronounced peaks at the cardinals compared to indoor scenes (Coppola et al., 1998; Straub and Rothkopf,2021) (see Supplementary Fig. 2). Thus we computed an average distribution across both indoor and outdoor scenes based on previous measurements (Coppola et al., 1998). We used this fixed prior distribution shown in Fig. 2a for all simulations and fits presented in this paper. Also, we consider four natural categories for orientation: vertical (‘V’), horizontal (‘H’), clockwise(‘CW’) or counterclockwise (‘CCW’) relative to vertical (Fig. 2b). We assume that there is some uncertainty associated with the categorical representation expressed in overlapping categorical distributions as well as due to noisy centers of the two cardinal categories that may vary trial by trial. Note that assuming a categorical structure that only distinguishes two categories (‘CW’ and ‘CCW’ relative to the vertical meridian) does not significantly change the model behavior (see Supplementary Figs. 3 and 4).

**Figure 2:**
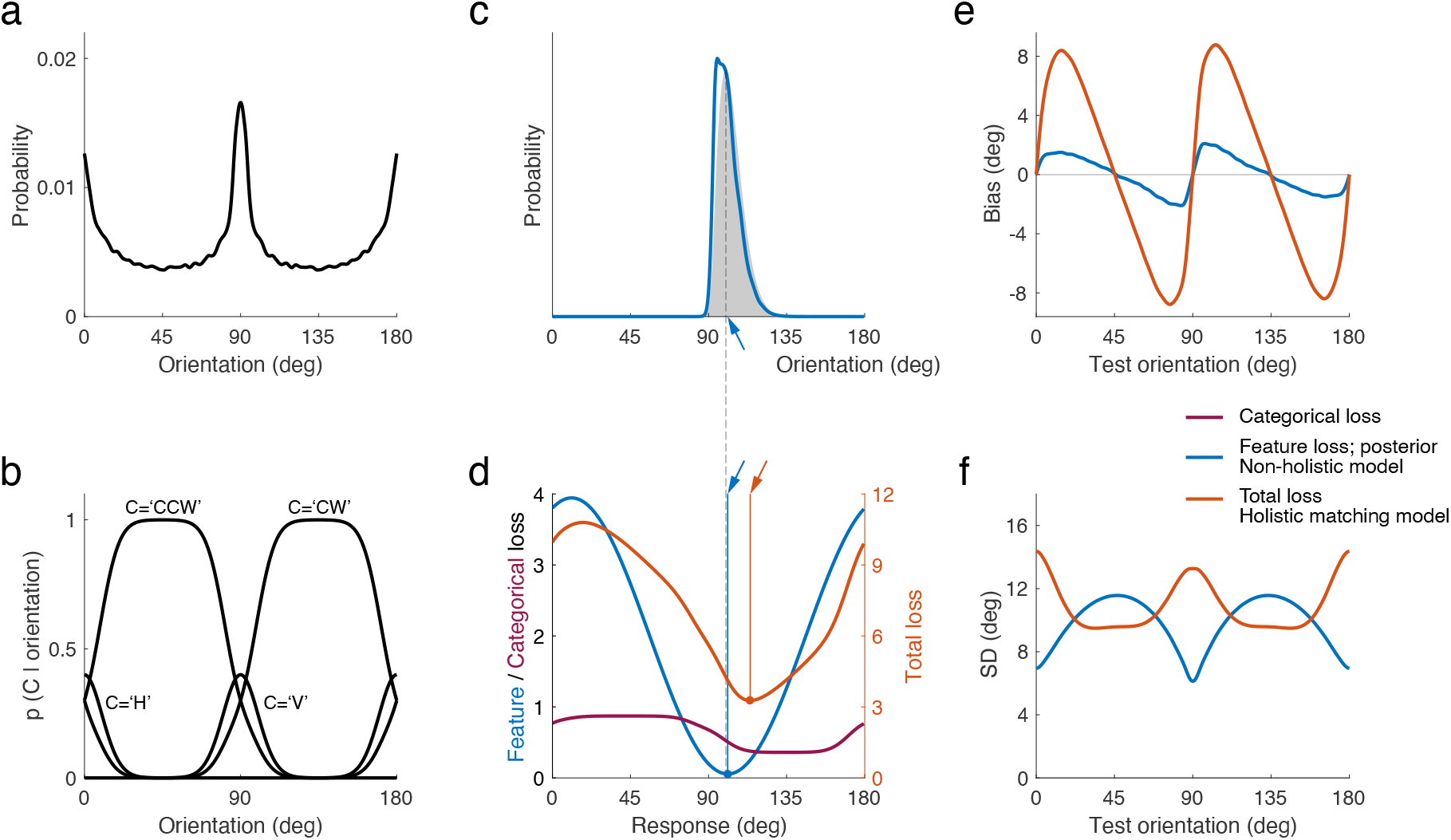
Model simulations for matching task with noiseless probe (typical condition). (a) Prior distribution used for model simulations and fits throughout this paper. It reflects the average statistics of local visual orientations in natural indoor and outdoor scenes measured by Coppola et al. (1998). (b) Category structure assumed by the holistic matching model. (c) Likelihood (shaded area) and posterior (blue curve) of test orientation for a given sensory measurement m (dashed line). The blue arrow marks the mean of the posterior, which is the optimal estimate predicted by the efficient Bayesian estimator. (d) Expected feature (blue), categorical (purple), and total loss (orange) given m (dashed line), and the optimal match predicted by the efficient Bayesian estimator (blue arrow) and the holistic matching model (orange arrow). (e) Bias pattern and (f) standard deviation in probe responses predicted by the two models. Supplementary Figure 2. Outdoor and indoor natural scene statistics.

Efficient encoding predicts likelihood functions that have long tails away from the nearest cardinal orientation for the assumed orientation prior (Fig. 2a). Figure 2c shows the likelihood function and the posterior distribution for a sensory measurement *m* of the test stimulus close to vertical (90 deg). Although the posterior is shifted towards vertical due to the prior, it inherits the long tail from the likelihood function. The long-tailed posterior distribution in combination with the loss function *L_θ_* is ultimately responsible for the predicted repulsive bias away from vertical of the efficient Bayesian estimator (Wei and Stocker, 2015). Figure 2d illustrates this by plotting the feature loss *L_θ_* for the same measurement m where the matching percept represents the point of minimal loss (arrow). This point of minimal loss, however, changes when considering the combined loss *L_tot_* of the holistic matching model. Because of category uncertainty, the minimum of the category loss *L_c_* does not coincide with the minimum of the feature loss *L_θ_*. Rather, it is shifted towards the center of the most probable category of the test stimulus resulting in larger repulsive biases (Fig. 2e). The precise bias pattern of the holistic matching model depends on the relative levels of feature (test and probe stimuli) and category uncertainty. Compared to the efficient Bayesian observer, the holistic matching model predicts larger repulsive biases for the same level of stimulus uncertainty.

### Matching experiment with noiseless probe stimulus: Data and model fit

We first validated the holistic matching model in the typical, noiseless probe condition using an extensive dataset from a previous orientation matching experiment (De Gardelle et al., 2010). In the experiment, subjects were asked to estimate the orientation of a briefly presented test stimulus by adjusting the orientation of a probe stimulus. Sensory noise of the test stimulus was modulated by varying the presentation duration (*Methods*). Figure 3 shows the full error distributions of the combined human subject data for each of the four presentation durations. The distributions exhibit the characteristic repulsive bias away from cardinal orientations, and show no apparent asymmetry between the two cardinal orientations. Bias and variability increase with decreasing stimulus presentation duration, which is a fundamental characteristic of Bayesian perception (Stocker and Simoncelli, 2006).

**Figure 3:**
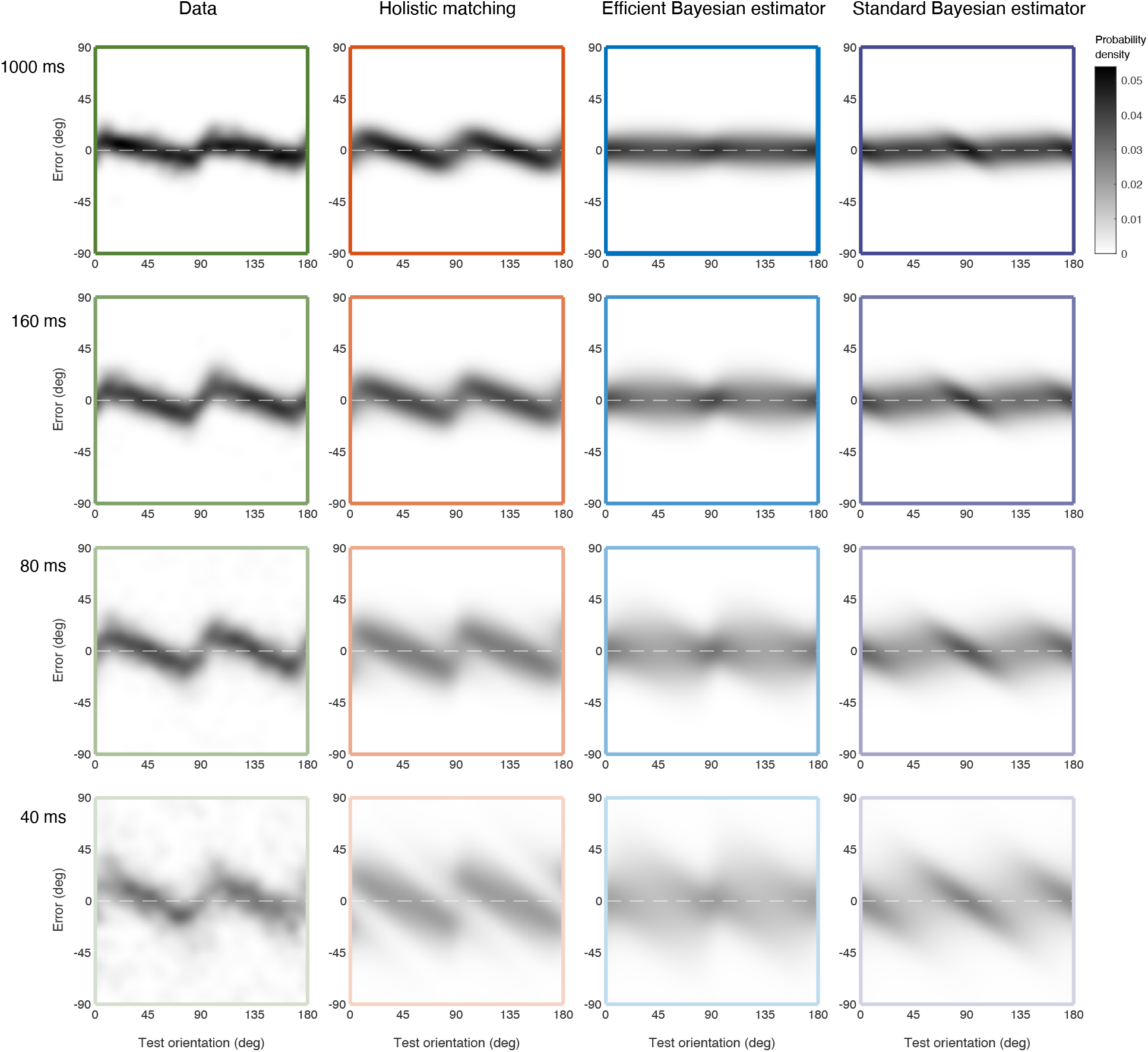
Data and model fits for matching task with noiseless probe. Shown are the error distributions of the matching responses for different presentation durations of the test stimulus (rows). Columns show the data (De Gardelle et al., 2010) and the corresponding best-fit model predictions, respectively. Data distributions show clear repulsive biases away from the cardinal orientations. Bias and variability increase with decreasing presentation duration. The overall pattern of the distribution is well captured by the holistic matching model across all conditions. While the efficient Bayesian estimator correctly predicts repulsive biases, the overall shape of the predicted error distributions does not match the data. The standard Bayesian estimator (homogeneous encoding) predicts attractive biases. See *Methods* for more details on the data and the models. Supplementary Table 1. Fit parameter values of the holistic matching model. Supplementary Figure 1. Fit category structure of the holistic matching model. Supplementary Figure 3. Model fit assuming two orientation categories.

We fit the holistic matching model as well as its non-holistic variant (i.e., the efficient Bayesian estimator) to the response distribution data using a maximum likelihood procedure. For comparison, we also included a standard Bayesian estimator with homogeneous sensory encoding (*Methods*). We assumed the probe stimulus to be noise-free as it consisted of a Gabor patch with one visible strip that was continuously present until subjects confirmed their choice. Note, that all models used the same formulation of the feature loss *L_θ_* and the natural orientation prior shown in Fig. 2a. The holistic matching model fully captures the entire shape of the error distributions for all noise conditions, which is not the case for the two Bayesian estimation models (Fig. 3). Their predicted error distributions are mostly centered around zero and show much larger variability for oblique than for cardinal orientations. Furthermore, as expected, the standard Bayesian estimator predicts attractive bias near cardinal orientations. In general, the estimation models exhibit distribution patterns that are substantially different from the data. The difference is also evident when comparing bias and standard deviation of the data with the predictions of the models (Fig. 4). Human subjects exhibit repulsive bias away from cardinal orientations with increasing amplitude for increasing sensory noise (i.e., shorter presentation duration), while the standard deviation of their response distribution is higher at cardinal compared to oblique orientations. Predictions of the standard Bayesian estimator are the exact opposite. Although the efficient Bayesian estimator qualitatively captures the repulsive bias pattern in the data and its dependency on sensory noise, the predicted overall bias magnitudes are too small. Like the standard Bayesian estimator, it also incorrectly predicts higher standard deviation at oblique compared to cardinal orientations. In contrast, the holistic matching model predicts bias and standard deviations that not only qualitatively but also quantitatively match the data. Note, that the predicted higher standard deviation at cardinal orientations is caused by the fact that for test orientations close to the categorical boundaries, small differences in sensory measurements across trials can lead to large differences in probe responses due to the categorical matching process. This additional, categorical bias offsets the increased sensory accuracy at cardinal orientations due to efficient coding.

**Figure 4:**
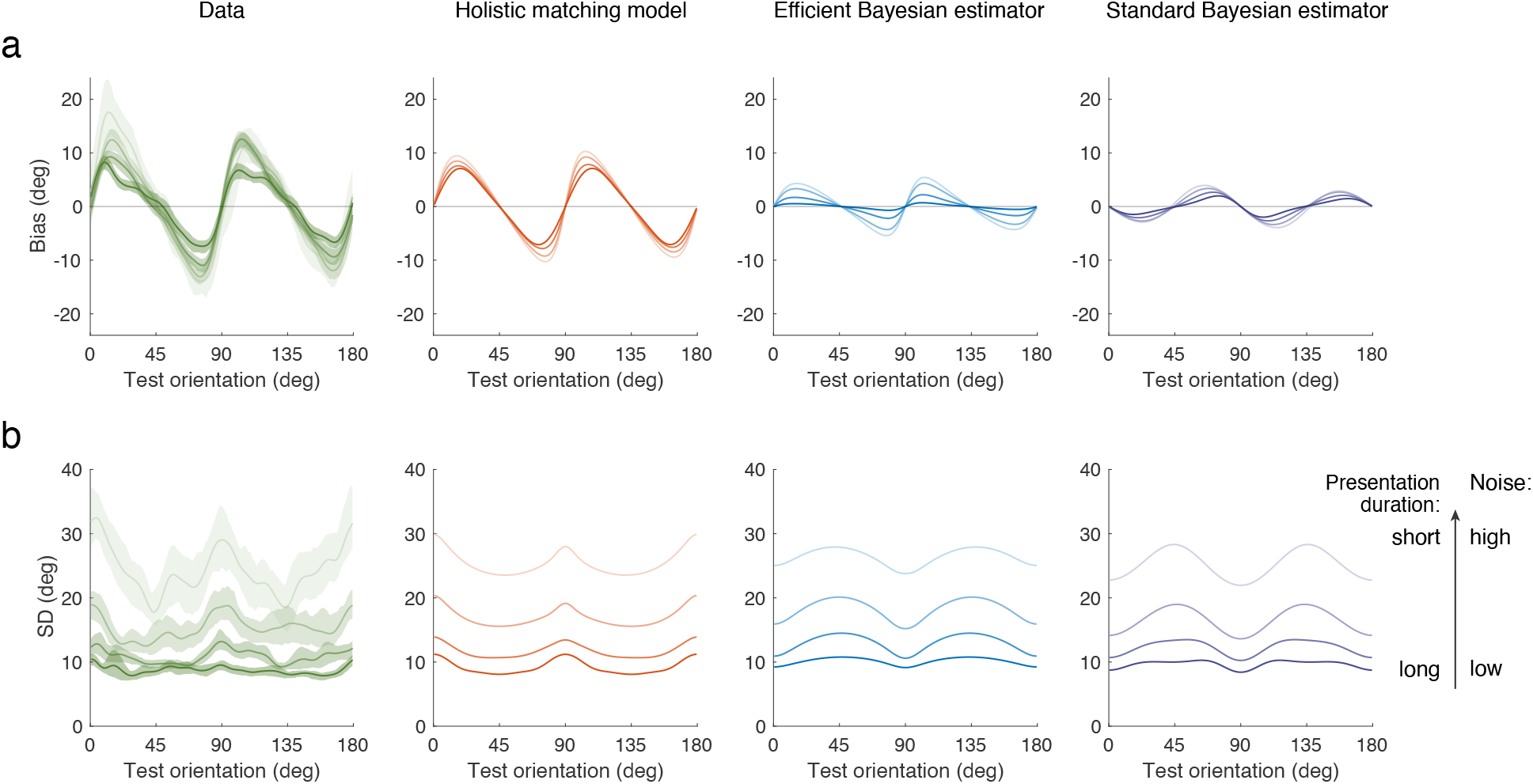
Data and model fits for matching task with noiseless probe: bias and standard deviation. (a) Subjects exhibit increasing repulsive bias with decreasing presentation duration. The holistic matching model fits both the pattern and the magnitude of the bias. The magnitude of the repulsive bias predicted by the efficient Bayesian model is too small compared to the data. In contrast, the standard Bayesian model predicts attractive bias. (b) Subjects’ variability is higher around cardinal orientations, which is well captured by the holistic matching model. The standard and the efficient Bayesian model predict the opposite. Data is re-analyzed from De Gardelle et al. (2010). Shaded areas represent 95% confidence intervals from 100 bootstrap runs. Supplementary Figure 2. Predictions of the holistic matching model using either the outdoor or the indoor scene statistics as the prior *p*(*θ*). Supplementary Figure 3. Model fit assuming two orientation categories.

We used cross-validation to quantitatively compare the performance of the models. We included an “omniscient” observer model in this comparison, which is an empirical model that directly transforms the training data distribution into a prediction probability of the error distribution using kernel density estimation. The omniscient observer serves as reference, indicating the best possible statistical prediction of the test set given the training set. As shown in Fig. 5, the holistic matching model predicts the data substantially better than the efficient and standard Bayesian model, and its performance is almost at the level of the omniscient model. This demonstrates that the holistic matching model provides an excellent account of orientation estimation behavior with a model complexity that does not lead to over-fitting of the data.

**Figure 5:**
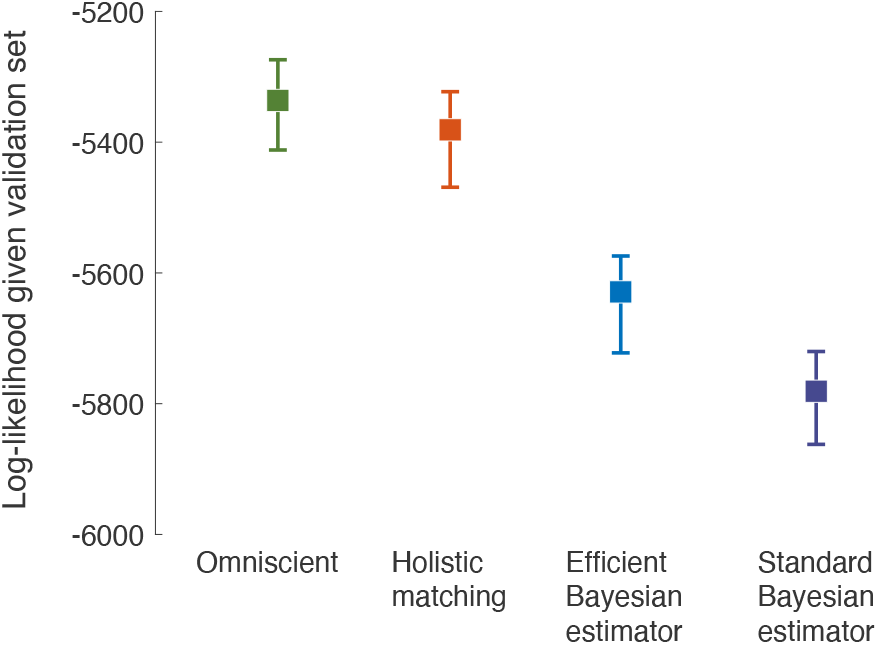
Cross-validation. Log-likelihood values of the model fit to the training set (80% of the data; randomly sampled), given the validation set (remaining 20% of the data). Squares represent the median and error bars indicate 95% confidence intervals over 100 repetitions. The holistic matching model performs significantly better than the efficient and the standard Bayesian observer model. The “omniscient” model is an empirical model that uses the data distribution in the training set as predictor of the validation data using kernel density estimation (*Methods*). The error bar of the holistic matching model and the omniscient model largely overlap, indicating that the holistic matching model explains the data as good as statistically possible. Supplementary Figure 6. Cross-validation of the omniscient model with different kernel sizes.

### Efficient sensory encoding is important for explaining human matching behavior

Is the categorical inference component of the holistic matching model sufficient to explain the repulsive bias pattern? To answer this question, we compared the predictions of the fit holistic matching model with and without efficient sensory coding. As illustrated in Fig. 6, without efficient sensory encoding (i.e., assuming homogeneous encoding) the model predicts decreasing bias magnitudes with increasing stimulus presentation time, which is opposite to the pattern seen in the data (Fig. 4a).

**Figure 6:**
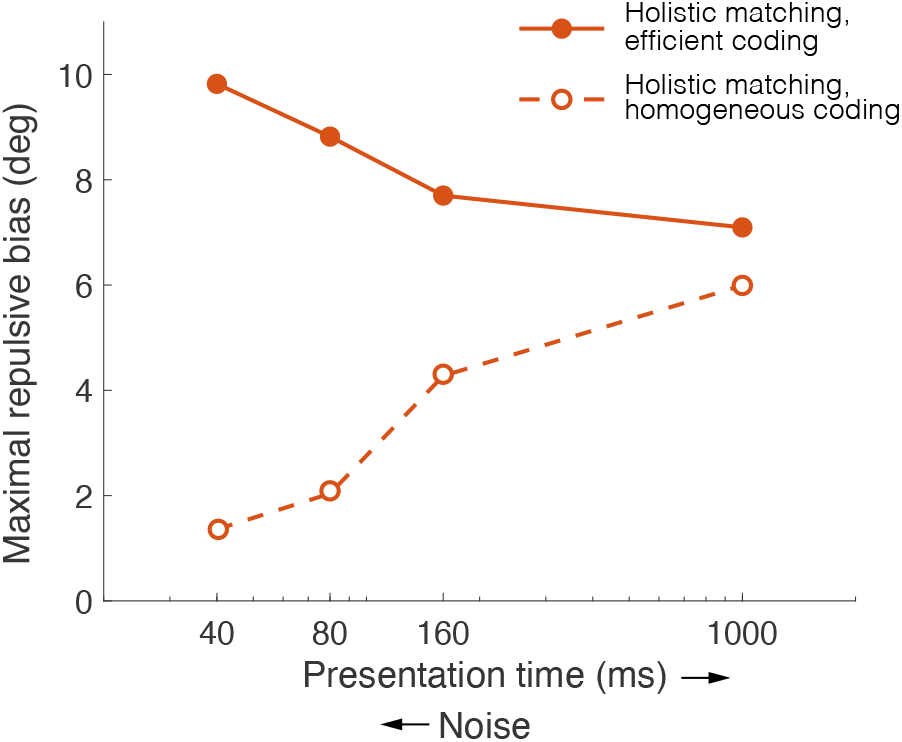
Effect of efficient sensory encoding. Maximum bias predicted by the holistic matching model with and without the efficient coding constraint. With efficient coding, the holistic matching model predicts decreasing bias magnitudes with increasing presentation times (i.e., decreasing sensory noise) consistent with the data. With homogeneous coding and all else equal, however, it predicts the opposite pattern.

Bias is modulated by sensory noise via three different processes of the model: sensory encoding, inference at the feature level, and inference at the categorical level. As sensory noise increases, the posterior distribution of the test orientation is more attracted to the peak of the prior distribution. At the same time, the difference in category posterior probability of the test stimulus decreases, leading to a flatter categorical loss curve (see Fig. 2d). Both effects lead to less repulsive bias as the sensory noise increases, which is the outcome shown in Figure 6 (dashed line). Efficient coding introduces repulsive biases by skewing the likelihood function away from the peak of the prior at cardinal orientations. However, larger sensory noise leads to more skewness in the likelihood function and therefore larger repulsive biases (Wei and Stocker, 2015). Thus, efficient coding is the one component of the holistic matching model that causes larger repulsive biases with higher sensory noise. As such, efficient sensory encoding is an indispensable assumption of the holistic matching model in order to accurately explain the data.

### Matching experiment with uncertain probe stimulus: Data and model fit

In most perceptual matching experiments the probe stimulus is unambiguous and noiseless, and thus its percept can be considered veridical. However, the holistic matching model explicitly models the perception of the probe orientation and thus can make predictions for more general uncertainty scenarios (Fig. 1b). Here, we specifically consider the case where stimulus uncertainty in test and probe are reversed.

Any model that only compares the feature estimates of the two stimuli would predict a reversal of the bias pattern when switching the role of the test and probe. However, the holistic matching model makes a qualitatively different prediction. Because the probe stimulus is adjusted to match the test also at the category level, adjusting the probe orientation towards the center of the most probably category of the test stimulus always reduces the expected categorical loss *L_c_*. This leads to a stable repulsive bias pattern whether the test and probe stimulus are interchanged or not. Figure 7 illustrates this qualitative difference between the model predictions.

**Figure 7:**
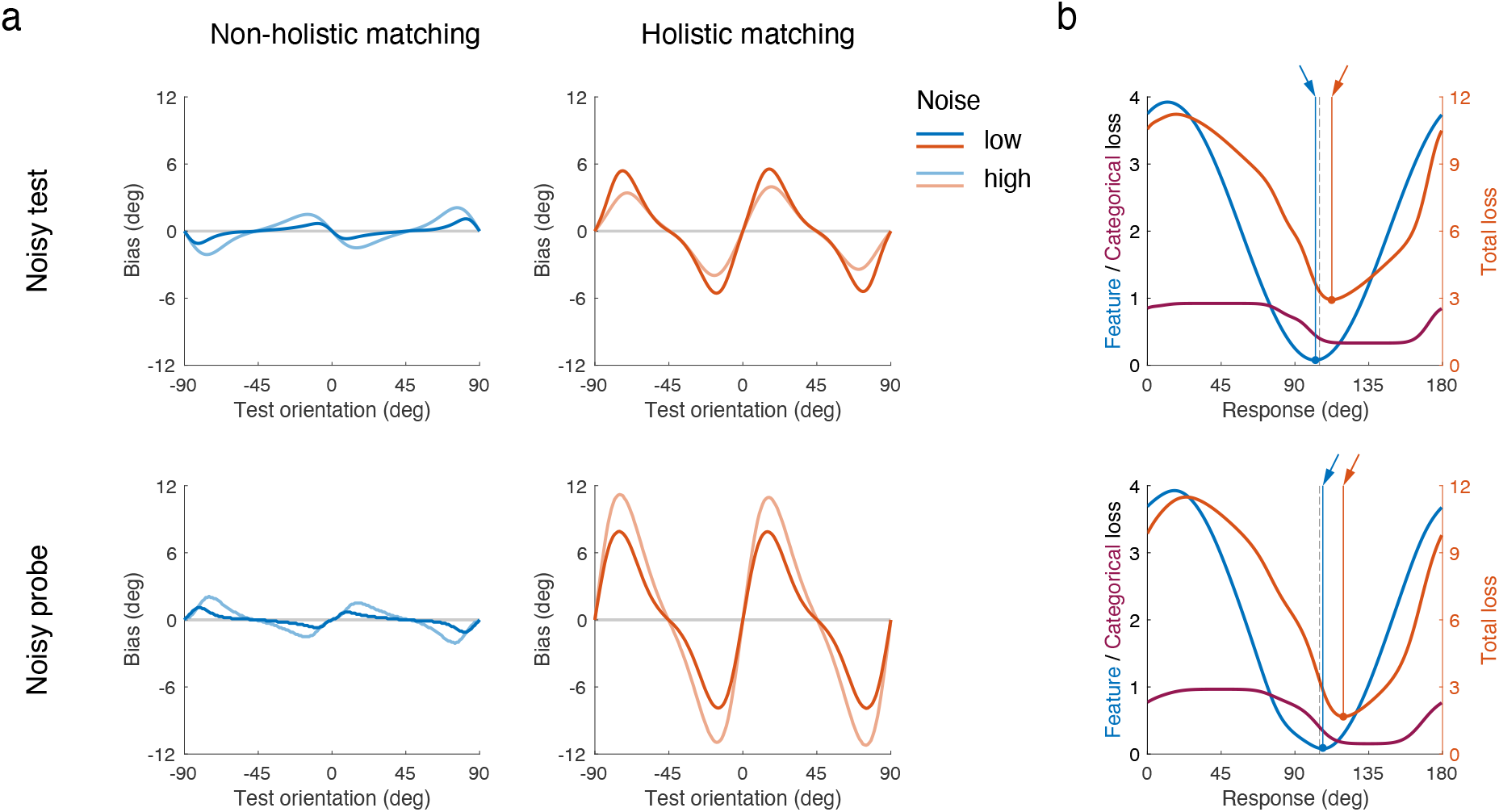
Model predictions for interchanging test and probe stimuli. (a) Estimation bias predicted by the holistic matching model (right) and its non-holistic variant (left) when the test has stimulus noise (top) or when the probe has stimulus noise (bottom). The non-holistic model variation is the full model shown in Fig. 1b yet without categorical inference; it is equivalent to the efficient Bayesian estimator if the probe stimulus is noise-free. The non-holistic matching model predicts opposite biases, while the holistic matching model predicts repulsive biases in both conditions. For the simulation we chose uncertainty to be mainly due to stimulus (external) noise and not sensory (internal) noise in order to qualitatively reflect the conditions of the experiment by Tomassini et al. (2010). Note, that this is the reason why the non-holistic model predicts attractive bias when the test has stimulus noise (see Wei and Stocker (2015)). (b) The models’ loss structures for a given set of sensory measurements. When the test and the probe stimulus are interchanged, the optimal response by the non-holistic model (blue arrow) flips to the other side of the measurement of the test (gray dashed line), but the optimal response according to the holistic matching model (orange arrow) remains on the same side because of the influence of the categorical loss.

The study by Tomassini et al. (2010) performed an orientation matching experiment where test and probe stimuli were interchanged. During the first half of the experiment, participants were shown an array of Gabor patches (test with stimulus noise) and were asked to adjust the orientation indicated by two dots (noiseless probe) to estimate the mean orientation of the Gabor patches. During the second half of the experiment, the roles of the stimuli were reversed; subjects were asked to rotate the array of Gabor patches (probe with stimulus noise) until the array orientation matched the orientation indicated by the two dots (noiseless test). Biases and standard deviations of subjects’ matching responses are shown in Fig. 8. As predicted by the holistic matching model, the biases are repulsive under both conditions and their amplitudes depend on the level of stimulus noise. We jointly fit the hierarchical model to the data across all conditions. The model well accounts for the observed repulsive biases in the small stimulus noise condition when the test stimulus is noisy and in the large stimulus noise condition when the probe stimulus is noisy. Likewise, when the test stimulus is noisy the predicted bias is close to zero for large stimulus noise but does not have a clear repulsive or attractive pattern, which matches the data (Fig. 8a). When the probe stimulus is noisy but the test stimulus is not, the bias is smaller in the small noise condition than in the large noise condition, which is the same pattern as in the data (Fig 8b). The standard deviation predicted by the model is for most part uniform with a magnitude that again is consistent with the data. The matching experiment by Tomassini et al. (2010) revealed human matching behavior that is well accounted for by the proposed holistic matching model, yet is difficult to even qualitatively reconcile with any estimation model.

**Figure 8:**
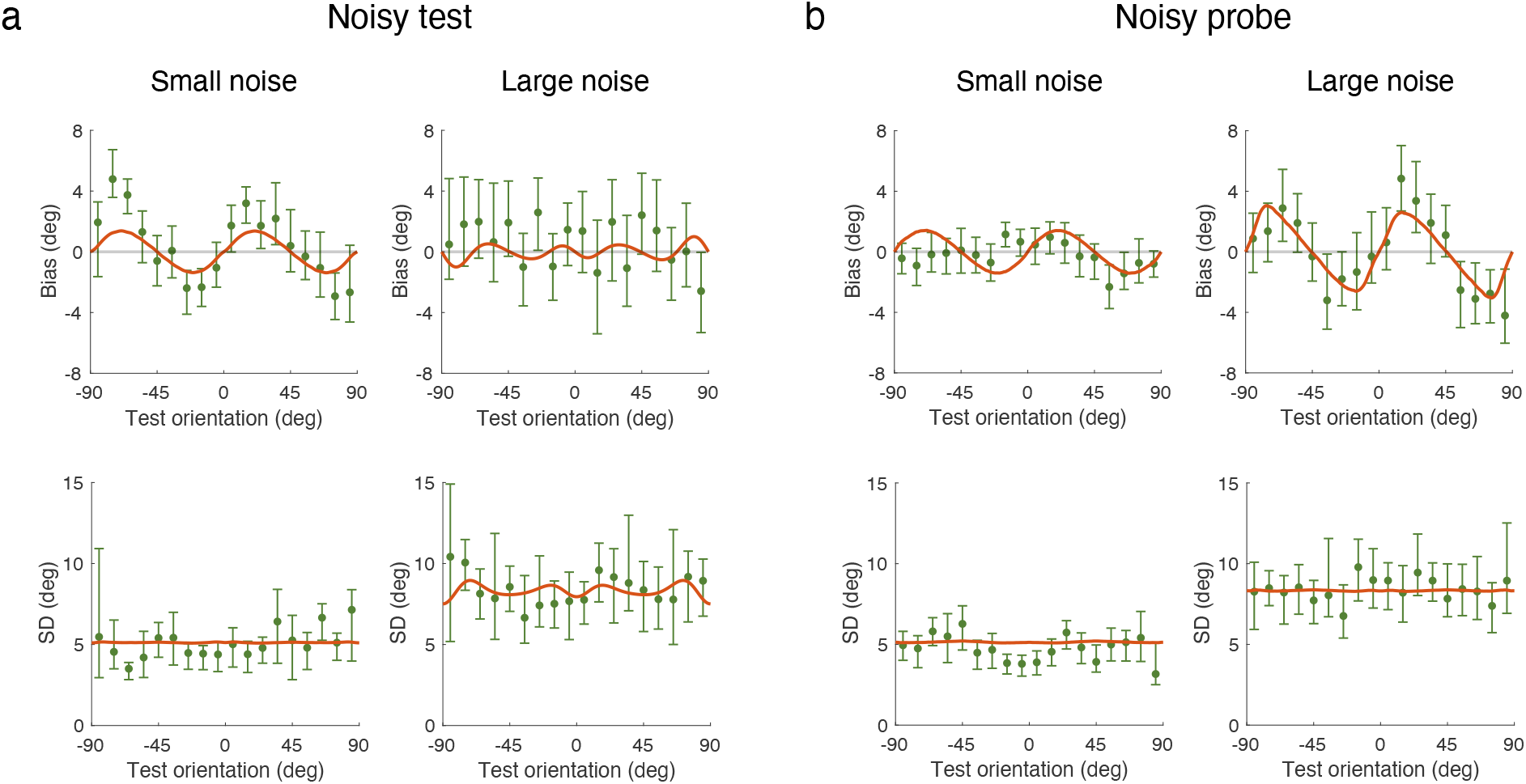
Data and model fits for interchanging test and probe stimuli for two different stimulus noise levels (combined subject). (a) Bias and standard deviation of subjects’ matching response data when the test stimulus is noisy and the probe is noiseless. (b) Same as (a) but probe and test stimuli are interchanged. The sign of the biases is not inverted. Biases are always repulsive or close to zero, depending on the level of stimulus noise. Solid lines represent the joint fit of the holistic matching model across all conditions. Data is re-analyzed from Tomassini et al. (2010). Error bars represent 95% confidence intervals from 100 bootstrap samples of the data. Supplementary Table 1. Fit parameter values of the holistic matching model. Supplementary Figure 1. Fit categories of the holistic matching model. Supplementary Figure 3. Model fit assuming two orientation categories.

### When to expect behavioral signatures of holistic inference

As the above data analysis and model comparison have shown, the holistic nature of the proposed matching model is responsible for increased (repulsive) response bias towards the category centers, different patterns in response variability, and the surprising finding that the bias pattern is not inverted when interchanging test and probe stimuli. However, the holistic matching model formally subsumes the efficient Bayesian estimator. Thus, we can clearly identify when we expect subjects’ behavior to show signatures of holistic inference, and when not because the models are equivalent.

The impact of the category level inference on behavior is determined by how gradually the expected categorical loss Lc decreases towards the center of the most probable category of the test stimulus (i.e., the slope of the categorical loss). Only if there is little category uncertainty and there is no uncertainty associated with the probe stimulus, then Lc approaches a step function with zero slope anywhere except at the boundary (Fig 9b, top left). In this case, the expected categorical loss is constant and independent of the probe orientation, and thus the expected response will be identical with or without holistic inference (Fig 9c, top). Otherwise, if there is category uncertainty (Fig. 9a, right) the probability of the probe category gradually changes with probe orientation, and thus the expected categorical loss also gradually decreases towards the center of the most likely category resulting in increased repulsive biases compared to the non-hierarchical model (Fig 9b and c, top). Likewise, if there is uncertainty associated with the probe stimulus then the expected category loss also gradually decreases for probe orientations towards the center of the most probably test category, which again results in increased repulsive biases (Fig 9b and c, bottom). We predict that signatures of holistic hierarchical inference are, to various degree, present in most psychophysical matching data. However, data from experiments that are designed such that they minimize subjects’ uncertainty about the category boundaries and use probe stimuli without uncertainty are expected to be equally well accounted for with the efficient Bayesian estimator.

**Figure 9:**
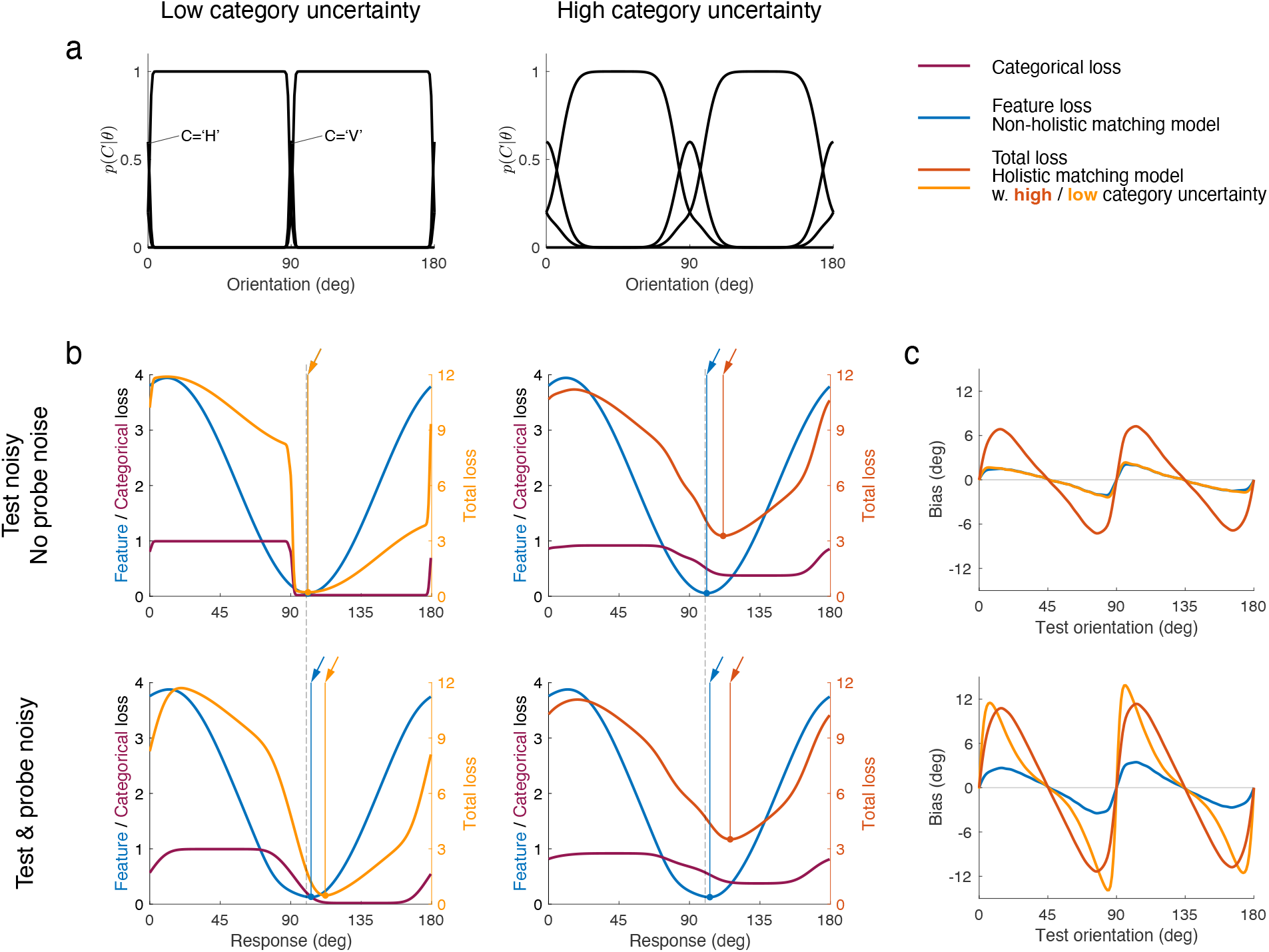
Model simulation with and without category uncertainty and with and without stimulus uncertainty in the probe stimulus. (a) Orientation categories *p*(*C*|*θ*) when there is low (left) or high (right) category uncertainty. (b) Expected loss for a certain measurement of test orientation (dashed line) when there is low (left column) or high category uncertainty (right column) and when the probe stimulus is (bottom row) or is not prone to stimulus uncertainty (top row). Arrows mark the optimal responses predicted by the non-holistic matching model (blue) or the holistic matching model (orange). (c) Bias predicted by the non-holistic (blue) and the holistic matching model when there is low (orange) or high (dark orange) category uncertainty, and when the probe stimulus is (bottom) or is not (top) prone to stimulus uncertainty.

## Discussion

We presented empirical and theoretical evidence that human sensory perception is a holistic inference process operating across a hierarchy of sensory representations. We introduced a holistic matching model to account for human behavior in a typical perceptual matching task for stimulus orientation. The model assumes that a subject’s response in this task represents an optimal match of the probe and the test stimulus in terms of a combined objective function considering both their differences in orientation and category identity. We validated the model against two different existing psychophysical datasets and demonstrated that, in addition to an efficient sensory encoding of the stimulus orientation, a holistic inference process is necessary in order to provide an accurate account of subjects’ full response distributions. The fact that subjects’ response bias is not inverted when switching the role of the test and the probe stimulus in the matching experiment is particularly strong evidence for the holistic matching model, because any model operating only at the feature level would predict the opposite behavior. The significance of our work lies in formulating and validating a novel, holistic inference theory for perception that explains why and how perception of a low-level visual feature (e.g., orientation) is dependent on its high-level structural representation.

Its holistic nature fundamentally separates our framework from existing Bayesian observer models that have been proposed to account for categorical effects in perception (Feldman et al., 2009; Bae et al., 2015; Kronrod et al., 2016; Landy et al., 2017; Bill et al., 2020). While these models share a similar generative process (Fig. 1b), inference in these models is limited to the feature level (i.e., orientation) by marginalizing over the entire generative hierarchy (i.e., categories). Marginalization effectively collapses the hierarchy, thereby reducing the inference process to a non-hierarchical Bayesian estimator with a heterogeneous prior determined by the weighted sum of the stimulus prior given each category. Thus the predictions of these models are identical to those of the non-holistic models considered in our model comparison (Fig. 3). Other studies have proposed that inference in these hierarchical Bayesian models is a sequential, top-down process where first the category of the stimulus is inferred before then computing the posterior at the feature level conditioned on the inferred category (Stocker and Simoncelli, 2007; Ding et al., 2017; Luu and Stocker,2018; Qiu et al., 2020) or the updated category belief (Lange et al., 2021), respectively. Although these “self-consistent” inference models predict increased perceptual biases away from categorical boundaries toward the center of the more likely stimulus category (i.e., confirmation biases), inference again is ultimately limited to an estimate at the feature level. Thus these models too can not explain why biases do not flip their sign when interchanging the probe and the test stimuli in a matching task(Tomassini et al. (2010); Fig. 8). Also, in contrast to these “self-consistent” inference models, the proposed holistic matching model is optimal and provides a rational, normative explanation for why and how categorical structures affect perceptual behavior. Sims et al. (2016) proposed a rate-distortion theory (RDT) based model using an objective function combining a cost at the feature and the category level similar to our approach. While the study showed how such optimal mapping can account for the estimation biases in color perception with regard to color categories, RDT is intrinsically an estimation model that would be rather difficult to adapt to modeling the matching process between test and probe stimulus under more general task conditions (i.e., with noisy probe stimuli). As a result, it too cannot account for the data by Tomassini et al. (2010).

It is worth highlighting the specific strengths of the proposed model, as well as its current limitations. First, our model makes detailed predictions of subjects’ behavior by specifying the entire response distributions, which permits a stringent and fine-grained model validation. This is in contrast to the many models that limit validation to some summary statistics such as the average response and its biases (e.g., Huttenlocher et al., 1991). Similarly, the model makes clear predictions in terms of meaningful parameters such as subject sensory noise levels, and the uncertainty and categorical structure of the stimulus. These are parameters that can be experimentally manipulated, allowing for a selective empirical tests of the model. Second, despite its complexity due to the hierarchical structure, the model is well-constrained. For all simulations and model fits, we used a fixed prior distribution over stimulus orientation given by measurements of the natural orientation statistics. This constraint likely prevents an even better quantitative account of the data yet demonstrates the robustness of our model (Supplementary Fig. 2). Furthermore, the model assumes that perceptual inference operates on efficient sensory representations of orientation, thus incorporating and extending previous work showing that human perception ubiquitously exhibits lawful hallmarks of efficient coding in combination with optimal Bayesian inference (Wei and Stocker, 2017). Thus, aside from the specification of the noise levels, the free model choices are essentially limited to the specification of the categorical structure of orientation. Somewhat surprising, there is little known about the natural categorical structure of human orientation perception. Thus our choice of ‘cardinal’ and ‘oblique’ discrimination reflects an intuitive assumption that, however, is shared with previous studies (e.g., Rosielle and Cooper, 2001). However, it is reassuring that assuming a categorical structure that only distinguishes between clockwise and counter-clockwise orientations across the vertical meridian does not significantly change the model behavior (see Supplementary Fig. 3 and 4 for the 2-category model fit to both datasets). Future experiments are necessary to better constrain the categorical representations of visual orientation in human observers.

Our model makes clear predictions when we expect signatures of holistic inference to emerge in human response behavior, and when not (Fig. 9). A recent study showed that data from three similar orientation matching experiments (Van den Berg et al., 2012; Bays, 2014; Pratte et al., 2017) can be well accounted for by the efficient Bayesian estimation model (Taylor and Bays, 2018). However, in line with our predictions, the differences in experimental design compared to the study by De Gardelle et al. (2010) explain why the Bayesian estimator was sufficient to fit the data. In particular, presenting the test stimulus at fixed equi-distant stimulus locations and using an unambiguous probe stimulus reduces both categorical and probe uncertainty, thereby reducing the influence of categorical inference on the matching response (Fig. 9). This is further supported by data of another recent study investigating the differences in orientation perception between individual with autism spectrum disorder and a neurotypical control group (Noel et al.,2021). In contrast to the just mentioned studies, this study used again a similar experimental design as in De Gardelle et al. (2010). Consistent with our interpretation, the study reports the same behavior signatures of holistic hierarchical inference as in De Gardelle et al. (2010), which is well accounted for by our holistic matching model (see Supplementary Fig. 5 for model fits). Future experiments that will systematically manipulate categorical uncertainty and noise in the probe stimulus will help to validate these predictions in more detail.

Finally, as low-level perception has been shown to follow common rules of sensory inference (e.g., Wei and Stocker, 2017), there is good reason to believe that our model generalizes to other stimulus domains. Color and speech perception have been shown to be categorical, with systematic reproduction biases towards the centers of the categories (Cibelli et al., 2016; Hardman et al.,2017; Chistovich, 1960; Kozhevnikov and Chistovich, 1966; Harris et al., 1961). Many perceptual domains that involve circular variables, such as motion direction (Rauber and Treue, 1998), pointing direction (Smyrnis et al., 2014), visual and vestibular heading direction (Cuturi and MacNeilage, 2013), exhibit repulsive bias away from cardinal directions just like orientation, and have also been shown to have better discrimination at cardinal directions (Gros et al., 1998). Similarly, studies in visuospatial memory distortion have found biases towards landmarks, which has been explained by the efficient Bayesian estimation model (Langlois et al., 2021). It will be interesting to investigate the degree to which a full quantitative account of these effects requires to consider not just efficient sensory representations but also a holistic hierarchical inference process as proposed here.

### Conclusions

Bayesian estimation models have been successful in accounting for many well-known distortions in perceptual behavior. In particular in combination with efficiency constraints on the sensory representations, they provide meaningful (normative) explanations for many of the characteristic bias and variability pattern observed in perceptual estimation tasks in terms of prior expectations and sensory uncertainty. Our results suggest, however, that it is time to augment these models to address the holistic nature of perception, where inferences at all levels of the representational hierarchy are combined to generate perceptual behavior even in simple low-level perceptual tasks. The novel, holistic matching model is a first step in this direction, providing a normative and intuitive explanation for how category representations affect perceptual behavior in a frequently used psychophyiscal task.

## Acknowledgments

We thank the members of the Computational Perception and Cognition laboratory for many fruitful discussions of this work. We thank Rafael Polania and Lingqi Zhang for helpful comments on the manuscript. This work has been supported in part by the National Science Foundation of the United States of America (Award IIS-1912232), and in part by the University of Pennsylvania. Preliminary results have been presented at the Annual Meeting of the Vision Science Society 2021.

## Methods

### Psychophysical data

#### Dataset by De Gardelle et al. (2010)

Each trial began with a background noise texture, then a test stimulus (Gabor patch) was presented for a variable duration at a random location 6.5 degs away from fixation. After the presentation of a mask and a blank interval, a randomly oriented probe stimulus (blue Gabor patch with only one visible strip) appeared at the test position. Participants were instructed to adjust the orientation of the probe using the mouse in order to reproduce the test orientation. Finally, they were also asked to report the visibility of the test stimulus on a continuous scale from 0 (“nothing seen”) to 1 (“fully visible”). Presentation times of the test Gabor were [1000, 160, 80, 40, 20] ms and 0 ms (no stimulus presented), randomly intermixed. 46 subjects participated in the experiment, divided into five groups. Four groups were presented with random test orientations in 2/3 of the trials and one particular orientation (vertical, horizontal, right or left oblique) in the remaining trials. The fifth group always received random test orientations. Each subject completed two to four blocks of 120 trials each. For our analysis, we combined the data of all five groups of participants but only included trials in which the test orientations were randomly selected. Furthermore, we excluded trials with presentation durations 20 ms (because data of those trials were too noisy to be reasonably analyzed) and 0 ms (because no test stimulus was shown). We also excluded trials for which the visibility rating was smaller than 0.01. After exclusion, the dataset contained 1103, 2187, 2140, and 1383 trials for each presentation duration, respectively.

Illustrations of the data distributions (Fig. 3) and bias and standard deviation (Fig. 4) are based on smoothing the raw trial data with a symmetric Gaussian kernel centered at each data point. Kernel size (standard deviation) was chosen to provide the most accurate density estimation based on cross-validation (5 degs; see Supplementary Fig. 6). Distributions are normalized to indicate the conditional probability of response for each test orientation. In order to allow for a fair visual comparison between models and data, we applied the same smoothing procedure for the model predictions shown in Figs. 3 and 4.

#### Dataset by Tomassini et al. (2010)

In the main experiment, subjects viewed an array of Gabor patches and adjusted the implied orientation of two dots, placed on opposite sides of the fixation mark, such that it matched the average orientation of the Gabor patches. In the control experiment, the test and probe stimuli were interchanged: subjects adjusted the orientation of the Gabor array to match the orientation indicated by the two dots. Adjustments were done by pressing two keys on the keyboard. The orientation of each Gabor patch in the array was randomly selected from a Gaussian distribution centered at the test orientation with two different standard deviations, resulting in two different stimulus noise conditions (Fig. 8). The orientation of the test stimulus was randomly selected from 18 orientations each 10 degs apart. For the main experiment, conditions with different fixed and response-terminated test presentation durations were measured in separate blocks. The control experiment only consisted of response-terminated presentations. Five subjects participated in the main experiment, each completing 8 trials per test orientation, presentation time, and stimulus noise level. Four subjects participated in the control experiment, each completing 16 trials per test orientation and stimulus noise level. Three subjects participated in both experiments. For our analysis, we combined the data across all subjects in both experiments, but only included the trials with response-terminated presentations. We also excluded trials in which the responses were 3 standard deviations away from the mean response.

### The holistic matching model

#### Efficient coding and feature inference

The following derivation follows Wei and Stocker (2015). Let *θ* be the orientation of the test stimulus and *m* its sensory measurement in a given trial. We assume that sensory encoding maximizes the mutual information between stimulus orientation and the sensory measurement (approximated by Fisher information (Wei and Stocker, 2016)), given that the total mutual information is limited. As a result, the prior distribution of the stimulus *p*(*θ*) and the Fisher information *J*(*θ*) of the sensory representation satisfy the efficient coding constraint

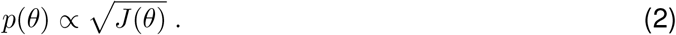

Sensory noise: consider a sensory space in which Fisher information is uniform. The optimal mapping 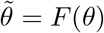 from stimulus to this sensory space is the cumulative of the stimulus distribution, thus *F*(*θ*) = ∫ *p*(*θ*)*dθ*. The likelihood function in stimulus space *p*(*m*|*θ*) can be computed by applying the inverse mapping 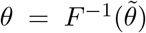 to the homogeneous likelihood function in sensory space 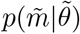 obtained by assuming uniform sensory noise according to a von Mises distribution

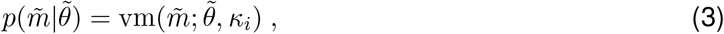

with *κ_i_* representing the sensory noise magnitude.

Stimulus noise: for the test stimulus in Tomassini et al. (2010) (array of Gabor patches) we assume uniform noise in stimulus space with stimulus samples *θ′* on each trial for a given stimulus *θ* following a von Mises distribution

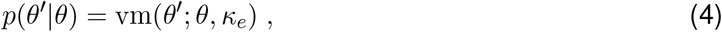

where *κ_e_* represents the stimulus noise magnitude. The stimulus sample *θ*′ corresponds to 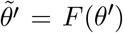 in sensory space and elicits a noisy sensory measurement 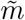 according to Eq. (3), hence

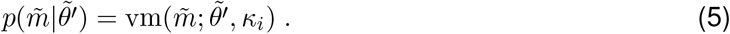

The distribution of the sensory measurement m in stimulus space is

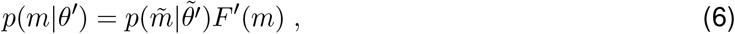

where 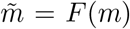. The likelihood function that takes both stimulus noise and sensory noise into account is

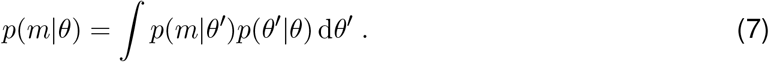

Finally, the posterior over stimulus orientation given the sensory measurement is

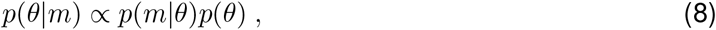

where *p*(*θ*) is the prior distribution over orientation, representing the natural orientation statistics (Fig. 2a).

#### Categorical inference

We assume four natural categories for orientation: vertical (‘V’), horizontal (‘H’), clockwise(‘CW’) or counter-clockwise (‘CCW’) oblique relative to vertical 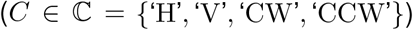. The horizontal category is defined by the von Mises distribution

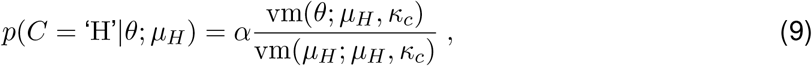

where *α* is the probability of the horizontal category at *μ_H_*, *κ_c_* represents the uncertainty in the categorical boundaries, and *μ_H_* represents a noisy signal of the horizontal orientation that may stochastically vary across trials according to

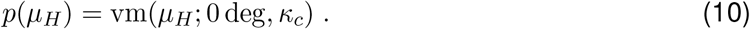

The vertical category is similarly defined with its center *μ_V_* always 90 degrees away from *μ_H_* (Eq. (10)). The oblique categories are the orientations in between the cardinal categories with a smooth transition given by the cumulative von Mises distributions

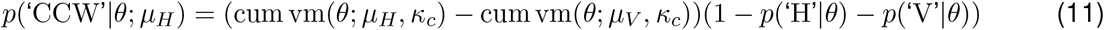

and

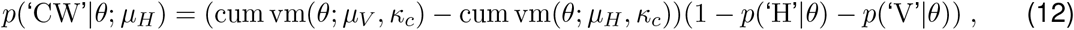

respectively. For simplicity, we assume a single parameter *κ_c_* to represent the uncertainty in the cardinal orientations and the uncertainty in the categorical boundaries.

Finally, the posterior distribution over category is computed as

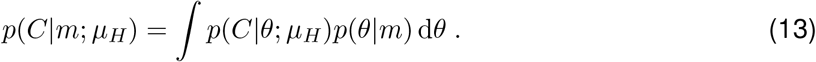

#### Matching

We assume that participants adjust the probe stimulus while obtaining continuous visual feedback about the probe orientation. Let *θ_p_* be the orientation of the probe, *m_p_* the sensory measurement of the probe, and *C_p_* the category of the probe. For simplicity, we assume that motor noise is additive, induced only after the probe is optimally adjusted (Fig. 1b). Note that we have considered more elaborate visuomotor control models that take motor noise into consideration when computing the optimal match, but found that the predictions do not significantly differ for the typical noise levels observed in orientation matching tasks.

The categorical loss is defined as whether the category of the probe is different from the category of the test orientation *C*:

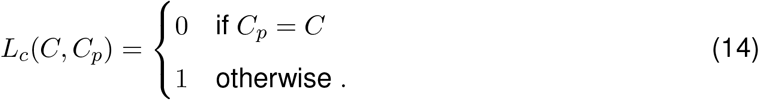

The feature loss is assumed to be the cosine of the difference between the probe and the test orientation, thus

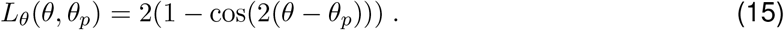

The cosine loss for circular variables is equivalent to the *L*_2_ loss for linear variables in the sense that the optimal estimate is defined by the (circular) mean of the posterior distribution.

The total loss (Eq.1) is the weighted sum of the feature loss and the categorical loss. Given the sensory measurements *m* and *m_p_*, the expected total loss is

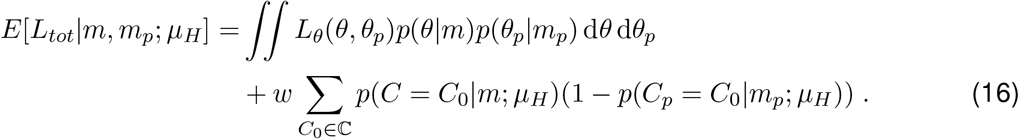

When there is no noise in the probe (*m_p_* = *θ_p_*) the expected total loss simplifies to

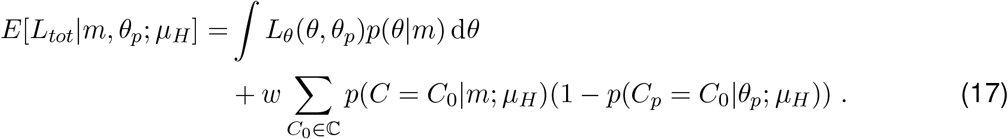

The optimal probe orientation 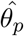 that minimizes the expected loss is

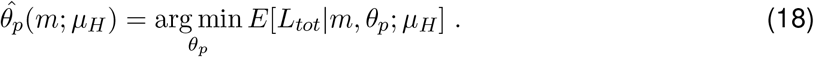

When there is noise in the probe, the observer has to minimize the expected loss based on the sensory measurement *m_p_*. For simplicity, we omit a description of how the observer adjusts the probe using visuomotor feedback. We simply assume that the observer adjusts the probe until they detect a probe measurement 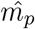 that minimizes the expected total loss, hence

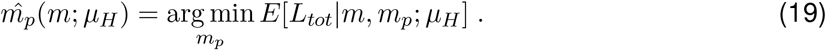

#### Predicted response distribution

Above is a description of the perceptual decision process of determining the optimal probe response, expressed by Eqs. (18) and (19), respectively. Now we calculate the predicted response distribution that follows from this process.

When there is noise in the perception of the probe stimulus, then there are different probe orientations 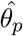 that could have generated the optimal probe measurement 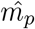. Since the probe orientation is generated by the observer and not the natural environment, we simply assume that the probability of the adjusted probe orientation given the optimal probe measurement 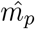 is proportional to the likelihood function as defined by Eq. (7) and is not affected by any non-uniform prior assumption, hence

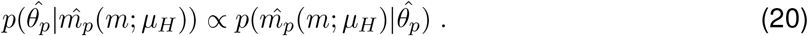

Because 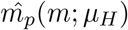 is a deterministic function (Eq. (19)) we can rewrite the probability distribution as

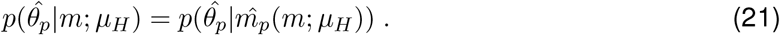

When the probe stimulus is noise-free, Eq. (21) turns into a Dirac delta distribution at the optimal 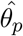 (Eq. (18)).

Finally, we assume that when the observer confirms the intended probe orientation *θ_p_* (e.g., with a button press), additive motor noise corrupts the answer leading to a noisy response 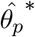 according to

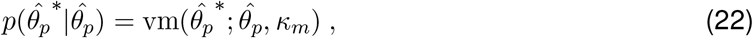

where *κ_m_* represents the motor noise magnitude.

Taken together, the predicted probability distribution of the matching response 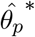 to a test orientation *θ* can be computed as

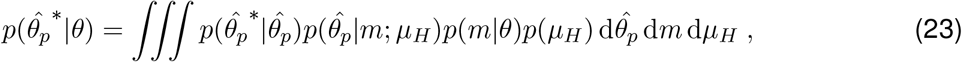

with the terms in the integral given by Eqs. (22), (21), (7), and (10), respectively.

### The non-holistic matching model

The non-holistic matching model shares the same feature inference process as the holistic matching model, but does not consider categorical inference (Fig. 1b). The matching process only consists of minimizing the feature mismatch between test *θ* and probe orientation *θ_p_*. The calculation of the response distributions for different noise conditions is also similar to calculations for the holistic matching model above.

If the probe stimulus is noiseless, the non-holistic matching model is equivalent to the *efficient Bayesian estimator* (Wei and Stocker, 2015) with the assumption that the probe orientation *θ_p_* is a direct representation of the optimal estimate 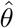 of the test orientation according to the loss function *L_θ_* (Eq. (15)) aside from potential, additive motor noise. The efficient Bayesian estimator shares the same efficient feature encoding as the holistic matching model described above.

The *standard Bayesian estimator*, however, assumes homogeneous encoding such that the sensory measurements m given the stimulus sample *θ*′ follows the von Mises distribution

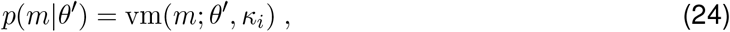

where *κ_i_* represents the constant sensory noise magnitude, independent of *θ*′.

### Model fitting

We jointly fit the model to the data of all the conditions in each dataset by maximizing the likelihood of the data given the model:

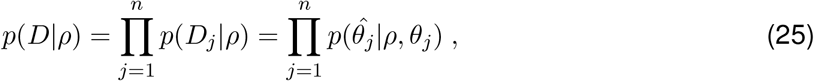

where *D* is the data, *ρ* represents the parameters of the model, *θ_j_* is the test orientation and 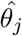 is the measured matching orientation (probe) in trial *j*, and *n* is the total number of trials.

We assume a fixed orientation prior for all model fits, representing the average natural orientation statistics extracted from indoor and outdoor scenes images (Coppola et al., 1998). More specifically, we apply a spline fit to the orientation histograms of indoor and outdoor scenes separately assuming that the distributions are symmetric around the vertical orientation (Supplementary Fig 2a), and then take the average of the two spline fits as the orientation prior (Fig 2a).

For fitting the data by De Gardelle et al. (2010), we assume no stimulus noise and four sensory noise levels corresponding to the four different presentation durations, resulting in a total of 8 free parameters:

- a group of four parameters *κ_i_* for four sensory noise levels;
- *κ_c_* for the uncertainty in the categorical boundaries;
- *α* for the probability of cardinal category;
- *w* for the weight of the categorical loss;
- *κ_m_* for motor noise.

For fitting the data by Tomassini et al. (2010), we assume one sensory noise level across all the conditions, and two stimulus noise levels corresponding to the two different standard deviations of the Gabor orientations in the stimulus array. So the holistic matching model fit contains 7 free parameters:

- *κ_i_* for sensory noise;
- a group of two parameters *κ_e_* for two stimulus noise levels;
- *κ_c_* for the uncertainty in the categorical boundaries;
- *α* for the for the probability of cardinal category;
- *w* for the weight of the categorical loss;
- *κ_m_* for motor noise.

The efficient Bayesian estimator has free parameters for sensory noise, stimulus noise, and motor noise; thus it has five free parameters for the data by De Gardelle et al. (2010), and four free parameters for the data by Tomassini et al. (2010).

The standard Bayesian estimator has the same free parameters as the efficient Bayesian estimator, except that for the comparison with the data by De Gardelle et al. (2010), we fixed the motor noise to be the same value obtained from the fit with the efficient Bayesian estimator (including the fit to the training set in each cross-validation run).

### Cross-validation

In each run of cross-validation, we randomly partition the data into a training set containing 80% of the trials and a validation set consisting of the remaining 20% of the trials. The partition is done separately for each noise level. We fit the model to the training set, then compute the likelihood of the fit model given the validation data. This likelihood represents the degree to which the fit model is supported by the validation data. We repeat this process 100 times.

#### The “omniscient” observer model

The omniscient model is an empirical model that serves as a reference for cross-validation. It directly considers the data in the training set as a prediction of the error distribution using kernel density estimation. Each data point in the training set is transformed into a symmetric 2D Gaussian probability kernel (diagonal covariance matrix). The resulting distribution is then normalized for each test orientation. The performance of the omniscient model on the validation set depends on the width of the Gaussian kernel: if the width is too small, the model over-fits the training set; if the width is too large, the prediction is too general and the model loses predictive power. We cross-validated the omniscient model with different standard deviations and found that a standard deviation of 5 degrees leads to the best performance (Supplementary Fig 6).

## Data and Code availability

All data in this paper have been previously published and were obtained directly from the authors. Matlab code for simulating and fitting the computational models will be deposited in the public repository Github upon the acceptance of the manuscript.

## Supplementary information

**Table 1:** Best-fitting model parameters for both datasets.

| parameter | value |
| --- | --- |
| **De Gardelle et al. (2010)** |  |
| $\kappa_i$: sensory noise | [356.94, 15.79, 4.58, 2.10] |
| $\kappa_c$: boundary noise | 8.29 |
| $\alpha$: cardinal probability | 0.60 |
| $w$: categorical weight | 10.75 |
| $\kappa_m$: motor noise | 34.63 |
| **Tomassini et al. (2010)** |  |
| $\kappa_i$: sensory noise | 698.50 |
| $\kappa_e$: stimulus noise | [651.39, 19.14] |
| $\kappa_c$: boundary noise | 4.79 |
| $\alpha$: cardinal probability | 0.52 |
| $w$: categorical weight | 0.52 |
| $\kappa_m$: motor noise | 38.20 |

**Table 2:** Best-fitting parameters of the 2-category holistic matching model for both datasets.

| parameter | value |
| --- | --- |
| **De Gardelle et al. (2010)** |  |
| $\kappa_i$: sensory noise | [211.47, 15.49, 4.59, 1.98] |
| $\kappa_c$: category noise | 58.61 |
| $\kappa_b$: boundary softness | 1.82 |
| $w$: categorical weight | 6.59 |
| $\kappa_m$: motor noise | 24.32 |
| **Tomassini et al. (2010)** |  |
| $\kappa_i$: sensory noise | 695.69 |
| $\kappa_e$: stimulus noise | [699.92, 17.56] |
| $\kappa_c$: category noise | 9.27 |
| $\kappa_b$: boundary softness | 2.33 |
| $w$: categorical weight | 0.69 |
| $\kappa_m$: motor noise | 37.29 |

**Table 3:** Best-fitting model parameters for data in the autism study by Noel et al. (2021).

| parameter | value |
| --- | --- |
| **Noel et al. (2021): Neurotypical subjects** |  |
| $\kappa_i$: sensory noise | 19.17 |
| $\kappa_c$: boundary noise | 8.25 |
| $\alpha$: cardinal probability | 0.56 |
| $w$: categorical weight | 6.98 |
| $\kappa_m$: motor noise | 27.56 |
| **Noel et al. (2021): ASD subjects** |  |
| $\kappa_i$: sensory noise | 6.19 |
| $\kappa_c$: boundary noise | 8.04 |
| $\alpha$: cardinal probability | 0.60 |
| $w$: categorical weight | 4.93 |
| $\kappa_m$: motor noise | 120.18 |

**Supplementary Figure 1:**
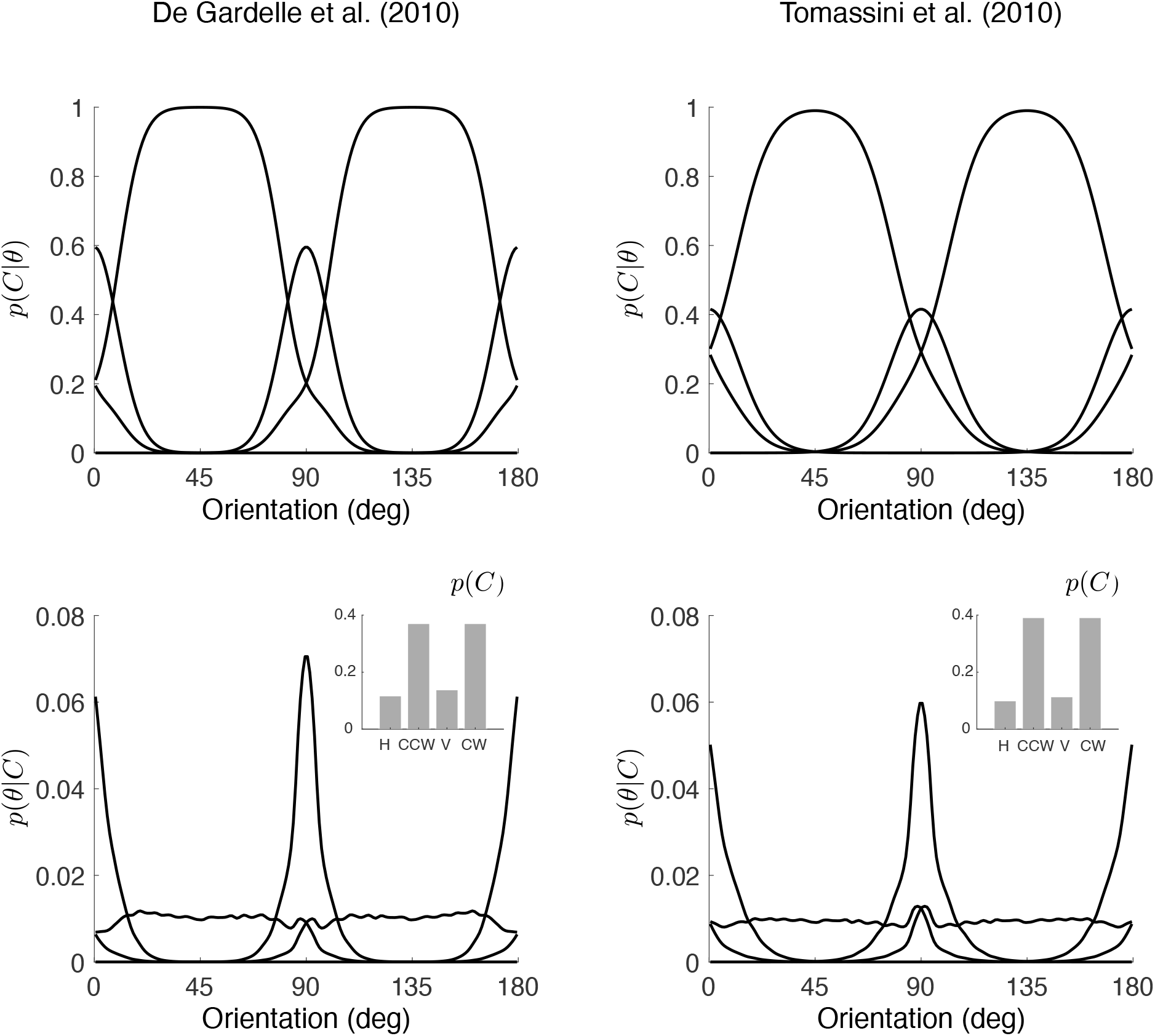
Best-fitting categories for both datasets.

**Supplementary Figure 2:**
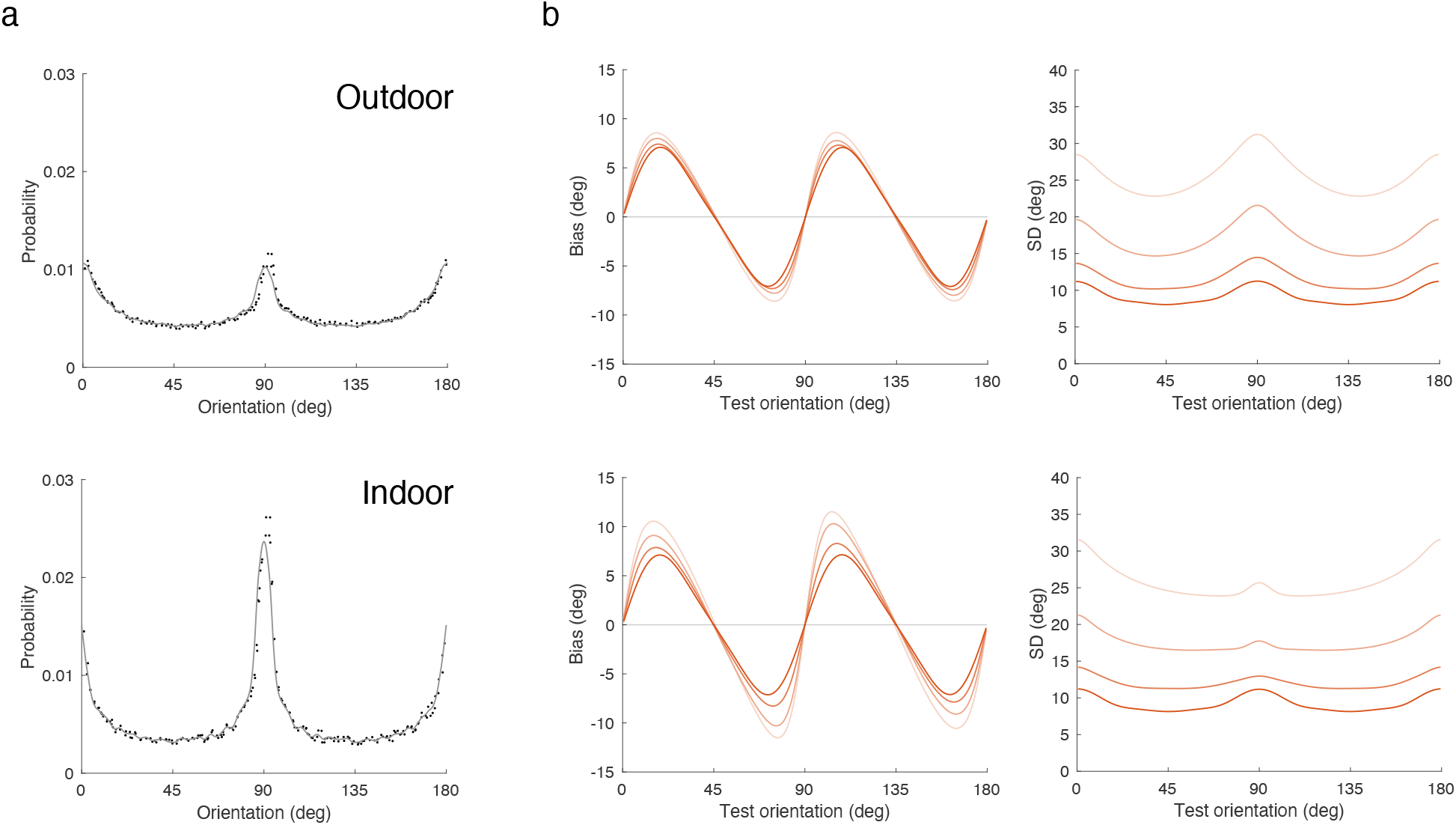
Simulations of the holistic matching model using “outdoor” and “indoor”orientation priors, respectively. (a) Image statistics of indoor and outdoor natural scenes (dots) and their smooth spline interpolations representing the corresponding prior distributions (lines). We assume the distributions to be symmetric around vertical. Data reanalyzed from Coppola et al. (1998). (b) Predicted bias and standard deviation of the holistic matching model using the two different prior distributions. All other model parameters where identical to the best-fit values listed in Table 1. Patterns in bias and standard deviation are qualitatively similar across the two priors. The peakier “indoor” prior leads to larger repulsive biases yet less pronounced differences in standard deviation compared to the “outdoor” prior. Simulations and model fits in the main text all use a fixed prior distribution that represents the average between the “indoor” and “outdoor” prior (Fig. 2).

**Supplementary Figure 3:**
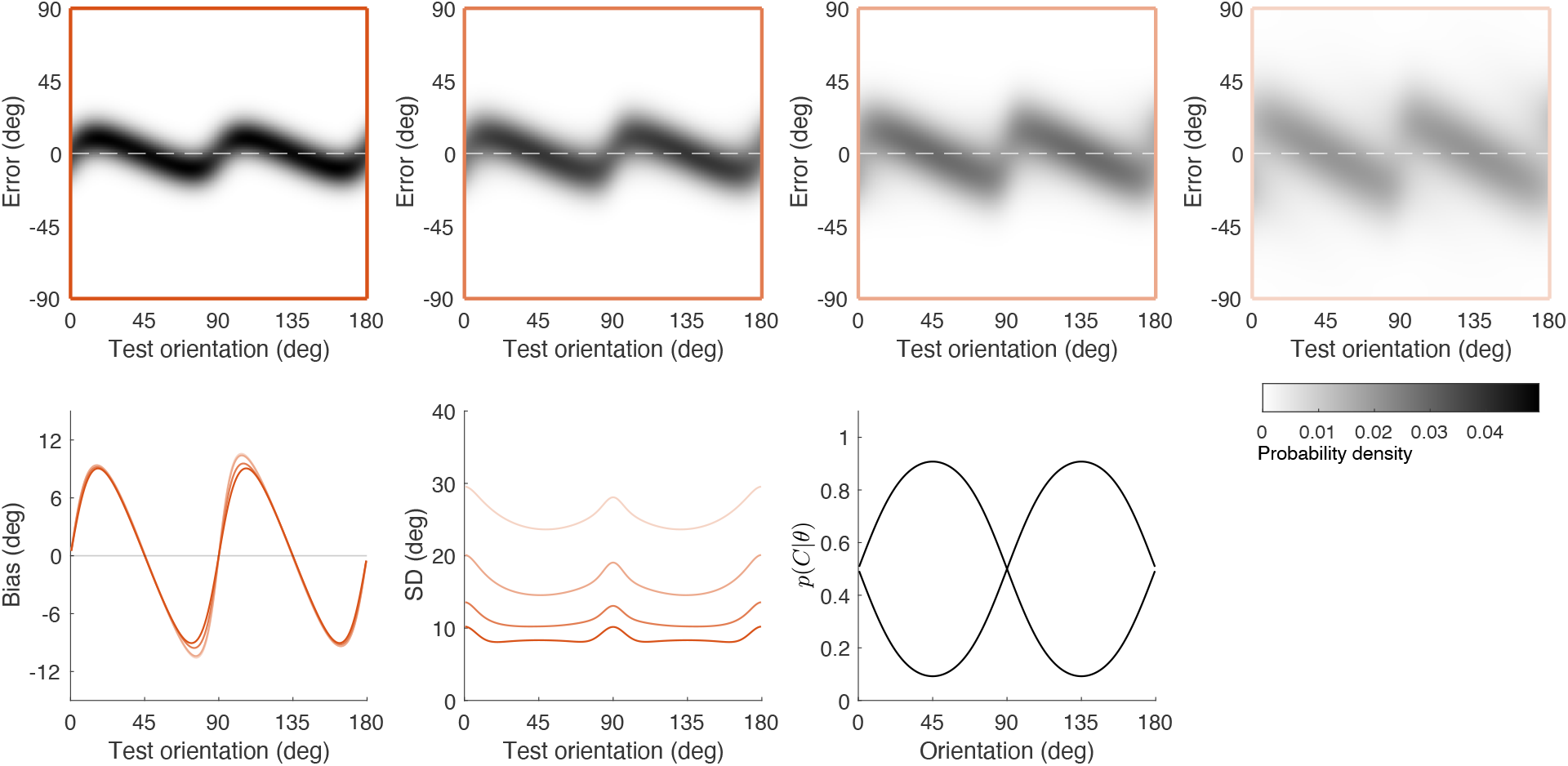
Two-category holistic matching model fit for matching task with noiseless probe. The cardinal probability *α* is set to zero, and the parameters for boundary softness and category noise are allowed to vary independently. The fitting procedure is otherwise identical to the model with four categories. Fit parameter values are listed in Table 2.

**Supplementary Figure 4:**
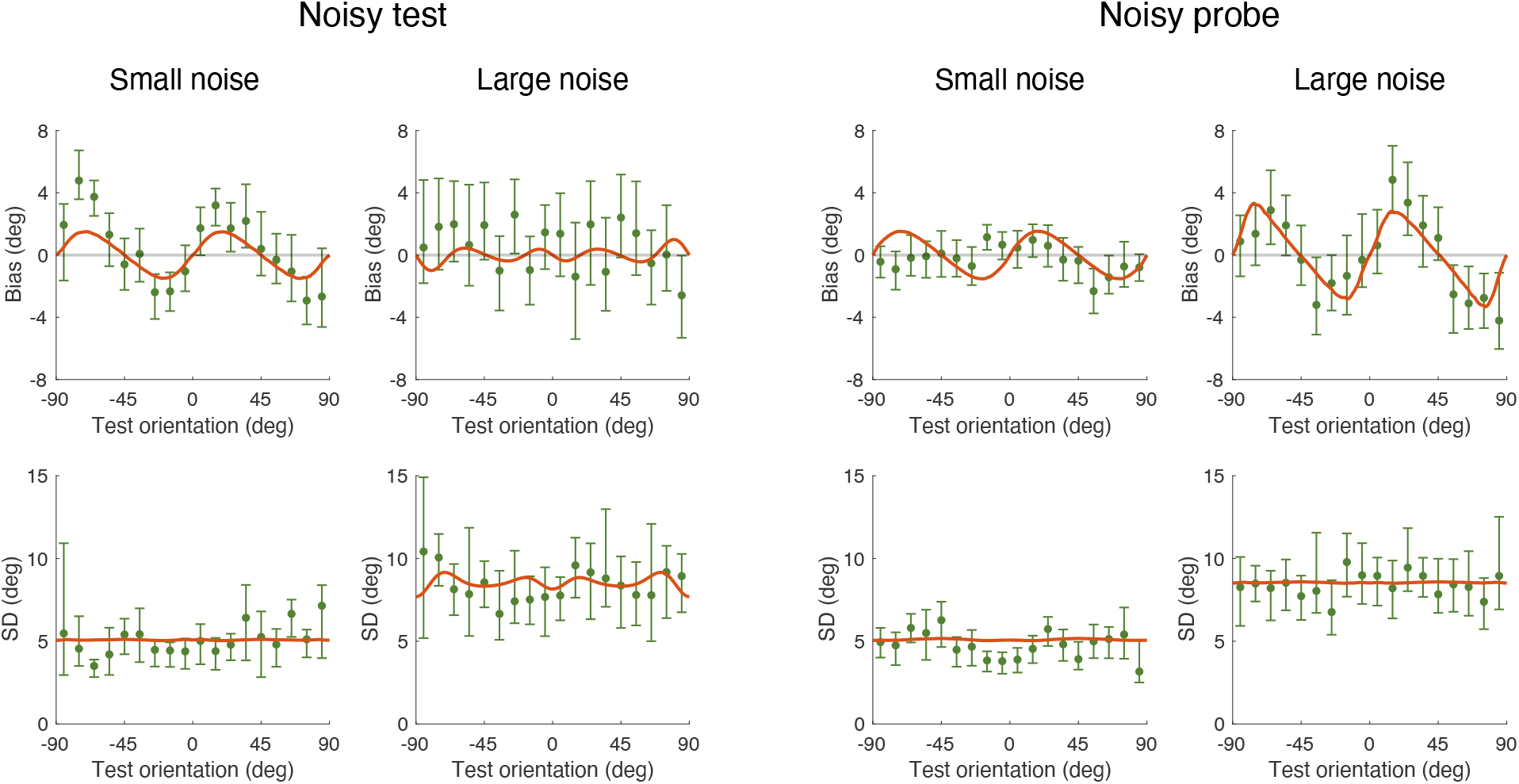
Two-category holistic matching model fit for matching task with noisy probe. The cardinal probability *α* is set to zero, and the parameters for boundary softness and category noise are allowed to vary independently. The fitting procedure is otherwise identical to the model with four categories. Fit parameter values are listed in Table 2.

**Supplementary Figure 5:**
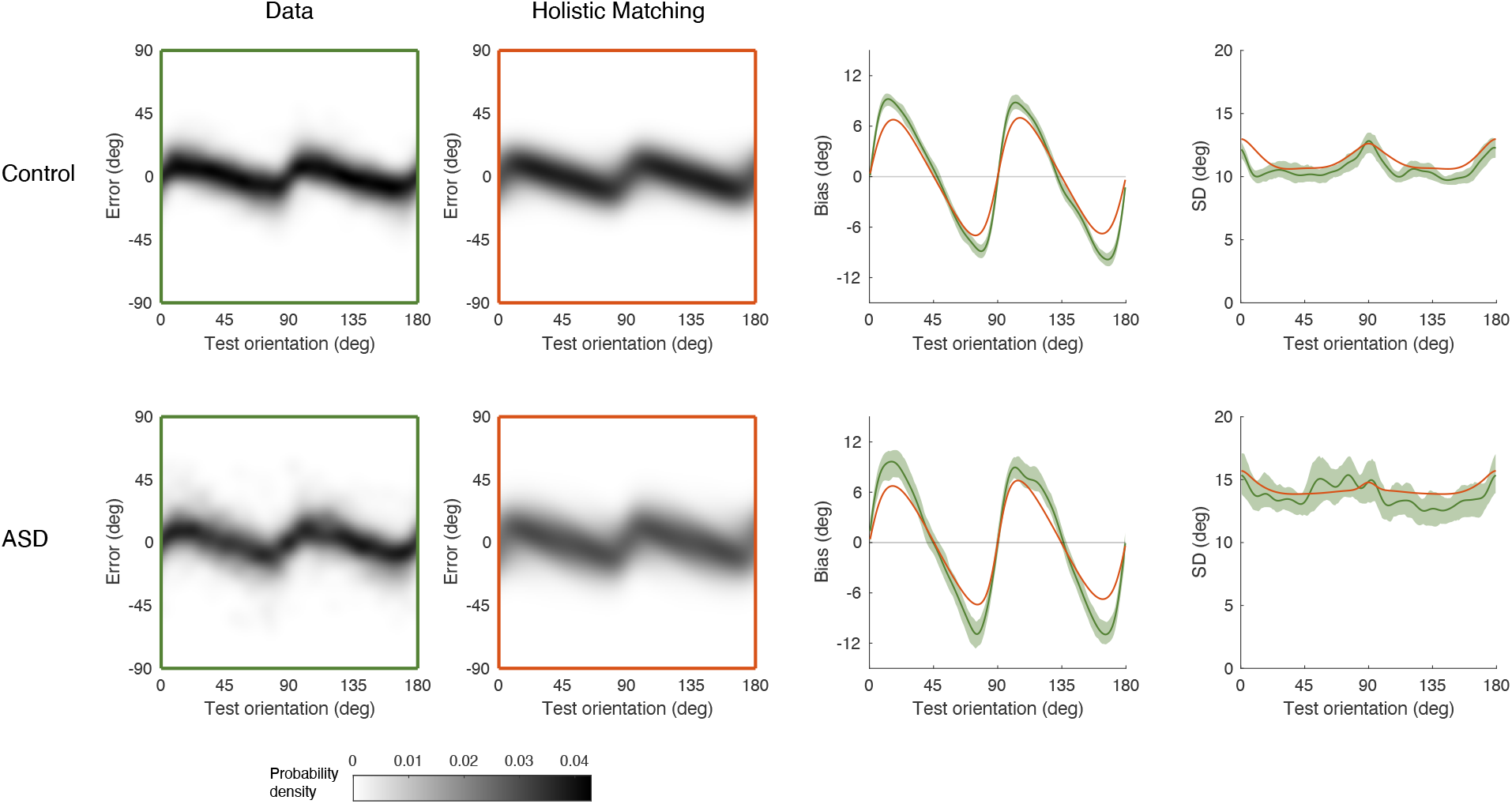
Data and model fits for matching task with noiseless probe in neurotypical and autistic (ASD) subjects. The data shows signatures for holistic matching, such as the shape of the error distribution, the relatively large bias magnitude, and the higher standard deviation at cardinal orientations than at oblique orientations. The data is well explained by the holistic matching model. Data are re-analyzed from Noel et al. (2021). Shown are data from only the no-feedback condition. Shaded areas represent 95% confidence intervals from 100 bootstrap runs. Fit parameter values are listed in Table 3. Note that the parameters are similar to those from the fit of the De Gardelle et al. (2010) data.

**Supplementary Figure 6:**
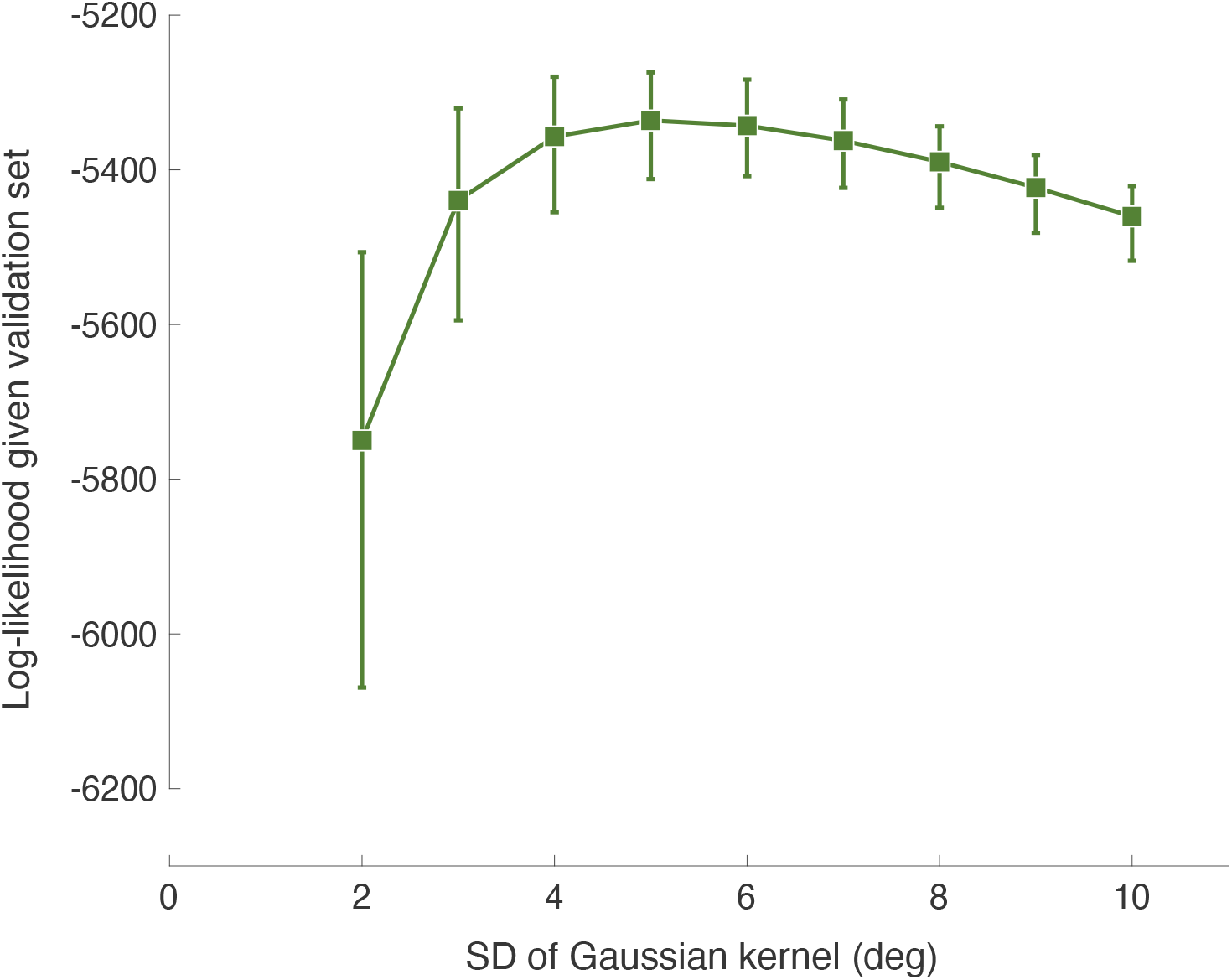
Cross-validation of the kernel density estimation accuracy for the omniscient model as a function of different Gaussian kernel size (standard deviation). Squares represent the median and error bars represent 95% confidence intervals of 100 repetitions of a repeated random sub-sampling cross-validation procedure. Accuracy shows a lawful dependency on kernel size with a standard deviation of 5 degrees providing the largest median likelihood value.

## Notes

### Competing Interest Statement

The authors have declared no competing interest.

